# Long-term NAD+ supplementation prevents the progression of age-related hearing loss in mice

**DOI:** 10.1101/2022.08.25.505332

**Authors:** Mustafa N. Okur, Risako Kimura, Burcin Duan Sahbaz, Uri Manor, Jaimin Patel, Leo Andrade, Kala Puligilla, Deborah L. Croteau, Vilhelm A. Bohr

## Abstract

Age-related hearing loss (ARHL) is the most common sensory disability associated with human aging. Yet, there are no approved measures for preventing or treating this debilitating condition. With its slow progression, continuous and safe approaches are critical for ARHL treatment. Nicotinamide Riboside (NR), a NAD+ precursor, is well tolerated even for long-term use and is already shown effective in various disease models including Alzheimer’s and Parkinson’s Disease. It has also been beneficial against noise induced hearing loss and in hearing loss associated with premature aging. However, its beneficial impact on ARHL is not known. Using two different wild-type mouse strains, we show that long-term NR administration prevents the progression of ARHL. Through transcriptomic and biochemical analysis, we find that NR administration restores age-associated reduction in cochlear NAD+ levels, upregulates biological pathways associated with synaptic transmission and PPAR signaling, and reduces the number of orphan ribbon synapses between afferent auditory neurons and inner hair cells. We also find that NR targets a novel pathway of lipid droplets in the cochlea by inducing the expression of CIDEC and PLIN1 proteins that are downstream of PPAR signaling and are key for lipid droplet growth. Taken together, our results demonstrate the therapeutic potential of NR treatment for ARHL and provide novel insights into its mechanism of action.

## INTRODUCTION

Age-related hearing loss (**ARHL**) is the most common sensory disability affecting the elderly human population. It manifests as a progressive, bilateral decline in hearing function starting with high-frequency sounds. One in three adults over the age of 65 show clinically diagnosable hearing loss, and the risk doubles with every decade of life *(1)*. Due to its slow progression, ARHL is often overlooked. Yet, it is extensively associated with cognitive decline, social isolation, and accident risk. Combined with the projected increase in the global elderly population, ARHL poses an enormous public health challenge.

Despite its rising prevalence and medical cost, the etiology of ARHL is not clear. Age-dependent changes in DNA damage accumulation, oxidative stress, mitochondrial dysfunction, and senescent-associated inflammation are postulated underlying biological mechanisms leading to ARHL *(2)*. Interestingly, the abundance of intracellular NAD+ levels plays a prominent role in correcting multiple aspects of the abovementioned biological events or ameliorating their cytotoxic outcomes *(3–5)*. NAD+ is a critical cofactor for numerous enzymes central to metabolism, longevity, and neuroprotection *(6)*. Notably, cochlear NAD+ levels decline in response to noise exposure, and NAD+ augmentation restores noise-induced neurite retraction from inner hair cells, suggesting that NAD+ levels are critical for proper cochlear function *(7)*. Indeed, we previously showed that NAD+ supplementation improves synaptic connectivity in the cochlea and prevents the progression of hearing loss in a premature aging disease model associated with dramatic hearing loss *(8)*. However, the impact of long-term NAD+ supplementation on ARHL has not been tested using a direct NAD+ precursor.

Nicotinamide Riboside (**NR**) is a NAD+ precursor found in foods including fruits, vegetables, meat, and milk *(9)*. Its oral intake is shown to boost NAD+ levels in a variety of tissues and organs including the brain and heart *(10, 11)*. NR is well tolerated in rodents and humans, providing a potential candidate for ARHL treatment *(12)*. Indeed, using two different mouse models, we show that long-term NR administration prevents the progression of ARHL, particularly of high-frequency sounds. Also, NR halts the further deterioration of already existing hearing loss in old mice. Mechanistically, we find that oral NR administration restores age-associated NAD+ decline in mouse cochlea. Also, our cochlear transcriptomic analysis demonstrates that NR upregulates pathways associated with synaptic connectivity. Our in-depth electrophysiological and histological analyses in cochlea support these results and show that NR enhances synaptic connectivity between cochlear sensory cells and afferent primary neurons in mice. In addition, we find that NR administration elevates the expression of key proteins involved in lipid droplet growth such as CIDEC and PLIN1, illustrating a novel pathway targeted by NR.

## RESULTS

### NR prevents the progression of ARHL

NAD+ levels decline upon aging in various rodent tissues including the kidney, brain, heart, and lung *(13)*. To assess whether the same phenomenon applies to cochlear tissue, we compared cellular NAD+ levels in the cochlea of young and old mice and tested if oral NR administration restores NAD+ levels. We found that total NAD+ and relative NAD+/NADH levels were indeed lower in the aged cochlea, and this decline was rescued significantly with NR administration (**Fig. 1a**). Given that NAD+ levels are associated with cochlear function *(7, 8)*, we next tested the effect of long-term NR administration on hearing loss in aged mice (**Fig. 1b**). One of the primary functions of NAD+ is to improve mitochondrial health by inducing mitochondrial turnover *(14, 15)*. To gain insight into the mechanism of action of NAD+’s benefit on hearing loss, we used WT (mtKeima) mice with a reporter gene to also assess mitochondrial degradation (mitophagy) in the cochlea *(16)*. We used the Auditory Brain Response (**ABR**) system to measure hearing capacity. ABR measures brain wave activity in response to sound stimuli at different frequencies (Hz) and decibel (dB) intensities (see methods for details). We observed an age-dependent elevation in hearing loss in the non-treated group at all frequencies tested, showing that the WT mouse strain used in these experiments suffers from ARHL (**Fig. 1c**). Remarkably, NR administration prevented the progression of ARHL specifically at high frequencies while no effect at lower frequencies was observed (**Fig. 1c**). This phenomenon was observed in both males and females (**Supp. Fig. 1**) with similar differences. We next compared the hearing threshold shift in treated and non-treated groups to analyze NR’s effect on hearing loss in individual mice. We found that NR not only prevented hearing loss progression but also improved high-frequency hearing in a subset of mice in the NR-treated group (**Fig. 1d**).

**Figure 1.**
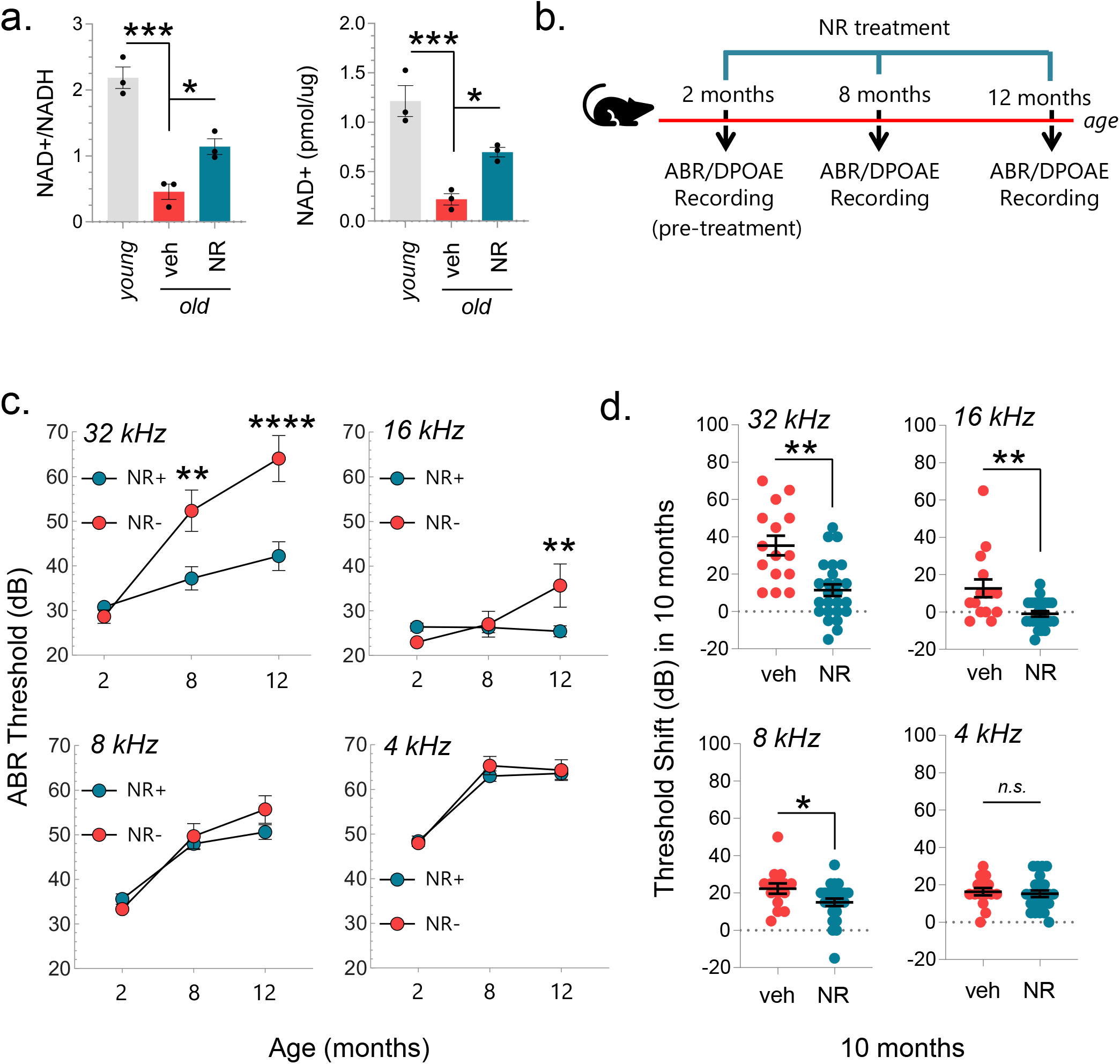
NAD+ supplementation using NR prevents the progression of age-related hearing loss in mice (mtKeima). **a**) The relative NAD+/NADH levels (left panel) and total NAD+ levels per mg of the cochlea (right panel) were measured in the cochlea of young (2-month-old), old (12-month-old), and NR-treated old mice (12-month-old). N = 3 and ordinary one-way ANOVA were used to determine significant differences. **b)** Outline for NR treatment and ABR/DPOAE recordings in mice. **c**) ABR thresholds for WT and NR-treated WT mice at 2, 8, and 12 m of age. A total of 40 WT mice were tested for hearing capacity at the age of 2 m and then randomly split into two groups of NR-treated (N=25) and non-treated (N=15). NR treatment started at the age of 2 m. ABRs in both groups were measured again at the age of 8 m and 12 m, which correspond to 6 m and 10 m of NR treatment respectively. Mixed effect analysis with Sidak’s multiple comparison test was used to determine significant differences. **d**) Threshold shifts at 8, 16, and 32 kHz in NR-treated and untreated groups. Note: ABR data in (**c**) were used to calculate the hearing shift. Two-tailed t-tests were used to determine significant differences. mean ± S.E., \**p* ≤ 0.05, \*\**p* ≤ 0.01, \*\*\**p* ≤ 0.001, \*\*\*\**p* ≤ 0.0001 and *n.s*., not significant.

### NR does not affect outer hair cell (OHC) function but elevates wave I and III amplitudes

Due to their vulnerable nature, OHCs are lost early during aging, prominently contributing to ARHL. Thus, we first tested if NR benefits hearing by preventing age-associated loss of OHCs. To address this, we measured DPOAE levels, which is an overall indicator of OHC health (see methods for details). We observed an age-associated decline in DPOAE levels in non-treated group at 32 kHz while no significant difference was observed at lower frequencies (**Fig. 2 and Supp. Fig. 2**). Interestingly, NR treatment did not impact overall DPOAE levels, suggesting that NR’s benefit on ARHL involves a mechanism other than the loss of OHCs (**Fig. 2**).

**Figure 2.**
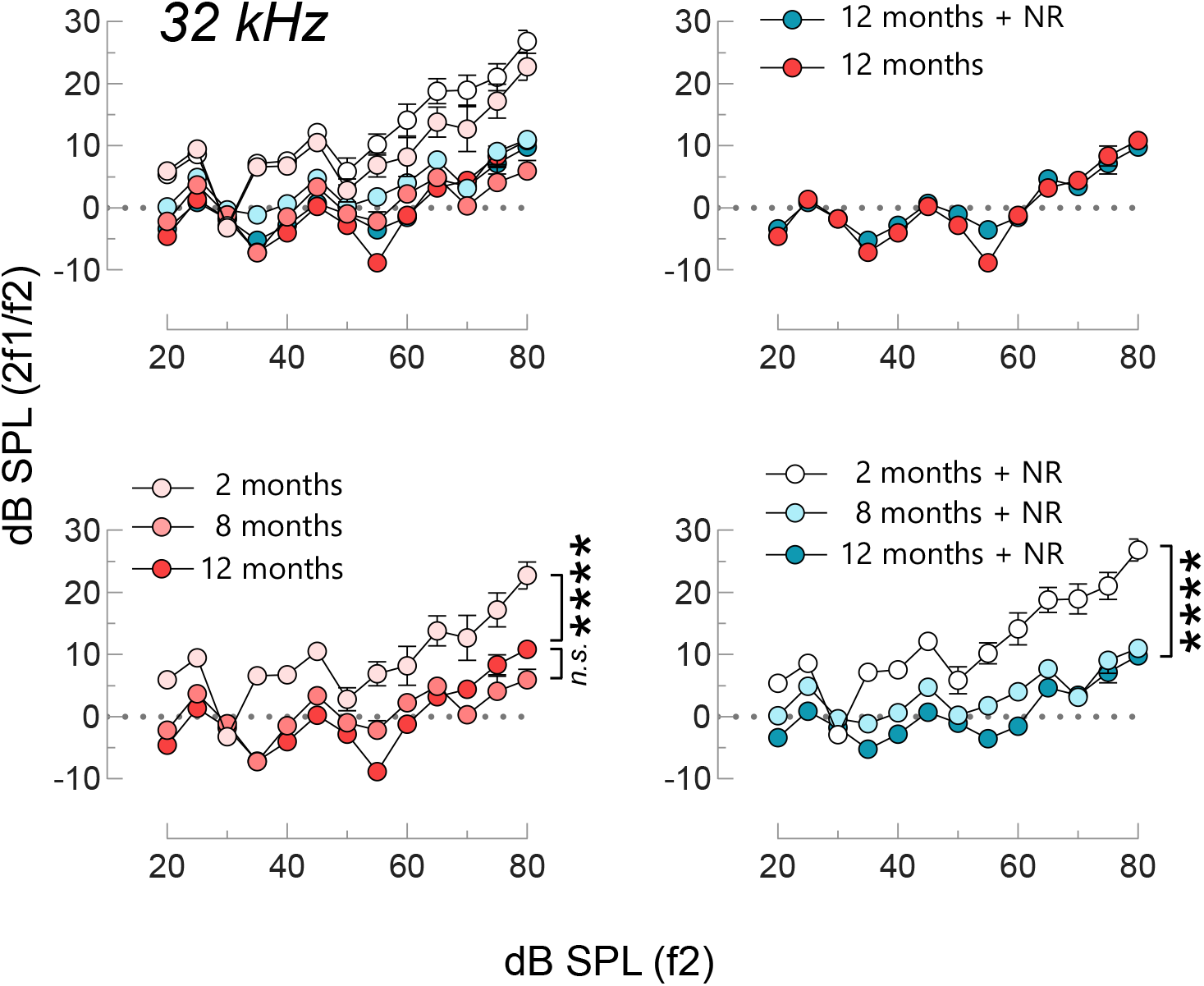
DPOAE levels at 32 kHz are shown at the age of 2, 8, and 12 m of age in NR-treated (N=25) and non-treated mice (N=13). The area under the curve (above −10 on the *y*-axis) is calculated for each sample and two-way ANOVA with Tukey’s post hoc test was used for statistical analysis. mean ± S.E., \**p* ≤ 0.05, \*\**p* ≤ 0.01, \*\*\**p* ≤ 0.001, \*\*\*\**p* ≤ 0.0001 and *n.s*., not significant.

ABRs are electrical potentials and recorded as five to seven waves in the first 10 ms due to synchronous firing of nerve fibers after an auditory stimulus. The first wave (wave I) captures the activity of inner hair cells (IHCs), afferent auditory neurons, and the synaptic connectivity between them. The following waves (waves II, III, IV, and V) are believed to correspond to locations descending through the auditory pathway corresponding to the cochlear nucleus, superior olivary complex, lateral lemniscus, and inferior colliculus, respectively. The reduction in the wave magnitude or increase in wave latency indicates a hearing deficit while the impacted wave provides valuable information for locating the deficit along the auditory pathway. We reconstructed average ABR waveforms and further examined the wave activity to gain more insight into NR’s benefit on hearing (**Fig. 3a)**. We found that the average wave amplitude (40 dB at 32 kHz) declines as a function of age in both treated and non-treated groups (**Fig. 3a**, compare the top and bottom panels). However, despite the decline, the amplitude of the waveforms in the NR-treated group was prominently higher than the ones in the non-treated group at 12 months (m) of age although they were similar at young ages (**Fig. 3a**, bottom panel). We next quantified and compared the amplitudes of individual waves in each mouse (**Fig. 3b**). Wave amplitudes fade away at lower decibel sounds so we analyzed wave amplitudes in response to higher decibel sound (80 dB) to account for all mice including the ones with prominent hearing loss. We found that NR specifically enhanced Wave I and Wave III amplitudes in the treated vs. non-treated group (**Fig. 3b**). These results indicate NR enhances cell function along the auditory pathway. This could include either or both sensory and auditory nerve cells and synapses thereof in the cochlea (Wave I), and nerve cells in the superior olivary complex in the brainstem (Wave III).

**Figure 3.**
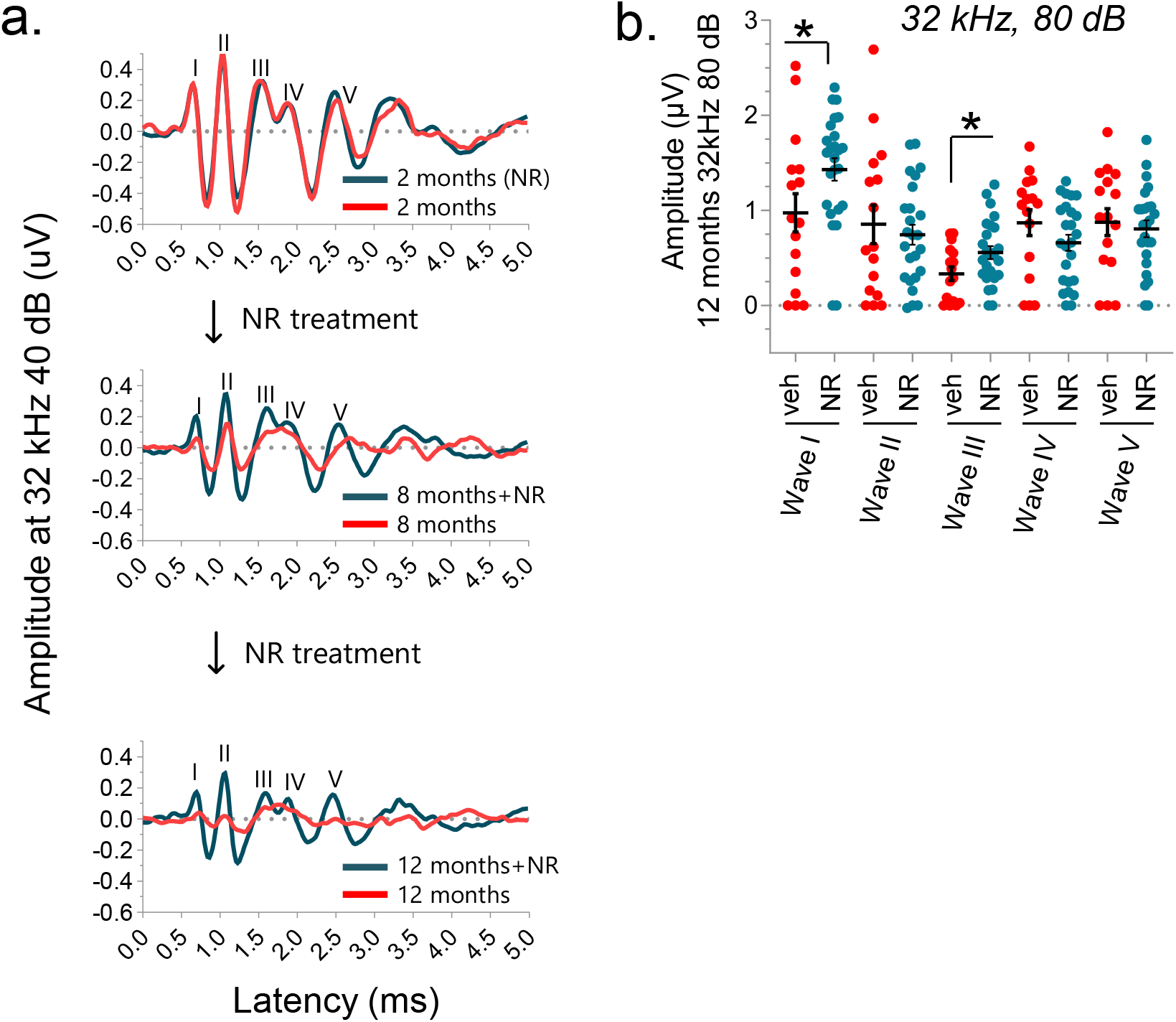
NR-treated mice show drastic preservation in waveform amplitude compared with age-matched controls at 8 and 12 m. **a)** Averaged ABR waveforms in NR-treated and non-treated WT mice resulting from a 32 kHz 40 dB SPL stimulus presented at 2, 8, and 12 m of age. **b)** Quantification of wave I-V amplitudes at 12 m. NR treatment significantly preserves wave I and III amplitudes. Each dot represents a single mouse. Two-tailed t-tests were used to determine significant differences. mean ± S.E., \**p* ≤ 0.05 and *n.s*., not significant.

### NR upregulates biological events associated with synaptic transmission and the PPAR signaling pathway

To investigate the biological mechanism of NR action on ARHL, we used unbiased RNA sequencing (RNA-seq) to identify the transcriptomic profiles of the cochlea from treated and non-treated groups. We found 1162 up-regulated genes and 1021 down-regulated genes with NR treatment (**Fig. 4a**). When hierarchical clustering was performed on those differentially expressed genes, NR-treated samples cluster more closely together, suggesting that NR led to a similar gene expression pattern in the cochlea in treated mice (**Fig. 4b**). We next performed gene ontology (GO) analysis to identify biological processes that are altered with NR treatment. Remarkably, NR treatment up-regulated many terms associated with synaptic transmission such as postsynaptic membrane, signal release from the synapse, synaptic vesicle transport, and synaptic vesicle cycling (**Fig. 4c**), consistent with the wave-form analysis in Figure 3, as well as with our previous study *(8)*. Down-regulated GO terms, on the other hand, included sensory perception, inner ear development, and ear development and morphogenesis. Notably, mitochondria-related terms were not among the most significantly changed GO-term list (**Fig. 4c**). This was surprising given the role of NAD+ in mitochondrial homeostasis. We extended our analysis and performed KEGG enrichment to identify associated biological pathways with treatment. We found that seven pathways were significantly up-regulated with treatment while no down-regulated pathways were observed (**Fig. 4d**). Interestingly, ‘synaptic vesicle cycle’ was an upregulated term in KEGG as it was in the GO analysis (**Fig. 4c and d**), supporting the notion that NR treatment promotes biological pathways associated with synaptic transmission. Among the upregulated list of KEGG pathways, we also identified that NR improved PPAR signaling, a family of transcription factors that regulate metabolic homeostasis *(18)*. Recent studies showed that PPAR activation protects the cochlea from oxidative stress, pointing out a potential biological pathway that NR targets *(19)*. PPAR regulates lipid metabolism and promotes the formation of lipid droplets, which are cellular organelles that store, release, and process lipids and proteins *(20)*. Remarkably, in the list of top genes that are most up- or down-regulated with treatment (**Fig. 4e**), we identified various genes such as *Cidec, Plin1, and Pck1* that are targeted by the PPARγ transcription factor and play key roles in lipid droplet formation. PLIN1 interacts with and activates CIDEC (FSP27) to regulate lipid droplet enlargement in mouse 3T3-L1 preadipocytes and human adipocytes *(21, 22)*. PCK1 is a metabolic enzyme that has a role in lipogenesis *(23)*. Although predominantly found in adipocytes, lipid droplets are present in most cells. Recent studies postulate that lipid droplets protect the delicate structure of the cochlea *(24)*. We therefore focused on CIDEC, PLIN1, PCK1, and PPARγ, validating their gene expression levels using RT-PCR. We also compared their levels to those in the young cochlea to determine changes with aging. Despite no significant effect of aging, the gene expression levels of *Cidec, Plin1*, and *Pck1* increased dramatically with NR treatment (**Fig. 5a**). These results were consistent with the gene expression patterns in RNA-Seq, confirming the reliability of RNA-seq results in this study. PPARγ levels, on the other hand, increased upon aging and showed no difference after NR treatment (**Fig. 5b**). We observed a similar trend in protein expression levels of these genes and found that NR significantly increases CIDEC levels while there was a trend towards elevated PLIN1 and PCK1 levels (**Fig. 5c-d**). Additionally, we used cultured mouse cochlear cell lines (HEI-OC1) to assess NR’s effect more directly. We found that only CIDEC’s levels were consistently elevated with NR treatment whereas there was no change in PCK1 levels and a slight reduction in PLIN1 levels in HEI-OC1 cells (**Fig. 5e-f**).

**Figure 4.**
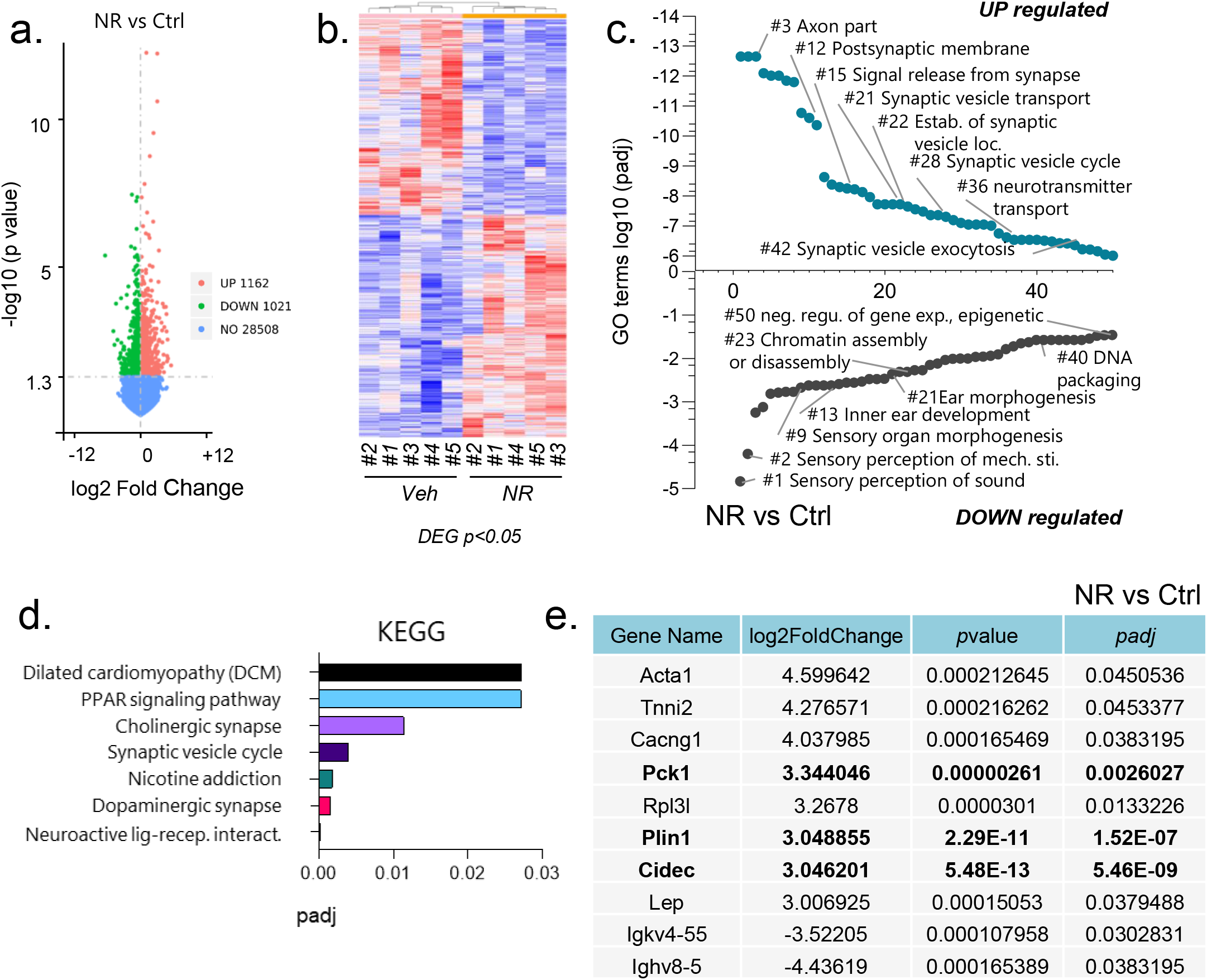
The transcriptomic analysis in NR-treated and non-treated WT mice cochlea. **a**) The number of up- and down-regulated genes with a *p*-value ≤ 0.05 in NR-treated and non-treated WT mice cochlea. **b**) Heatmap showing clustering of differentially expressed genes (DEG) (*p*-value ≤ 0.05) in NR-treated and non-treated WT mice cochlea. **c**) Graph showing the top 50 up- or down-regulated GO terms from the WT cochlea ±NR treatment. A padj-value ≤0.05 were the cutoff used for significance. **d**) Graph showing the up-regulated KEGG pathways from the WT cochlea ±NR treatment. A padj-value ≤0.05 were the cutoff used for significance. No significant down-regulated KEGG pathways were detected. **e**) The table shows a list of the top 10 genes with the highest value of fold change (up- or down-regulated). A padj-value ≤0.05 and fold-change (log2) ≥3 were the cut-offs used for significance.

**Figure 5.**
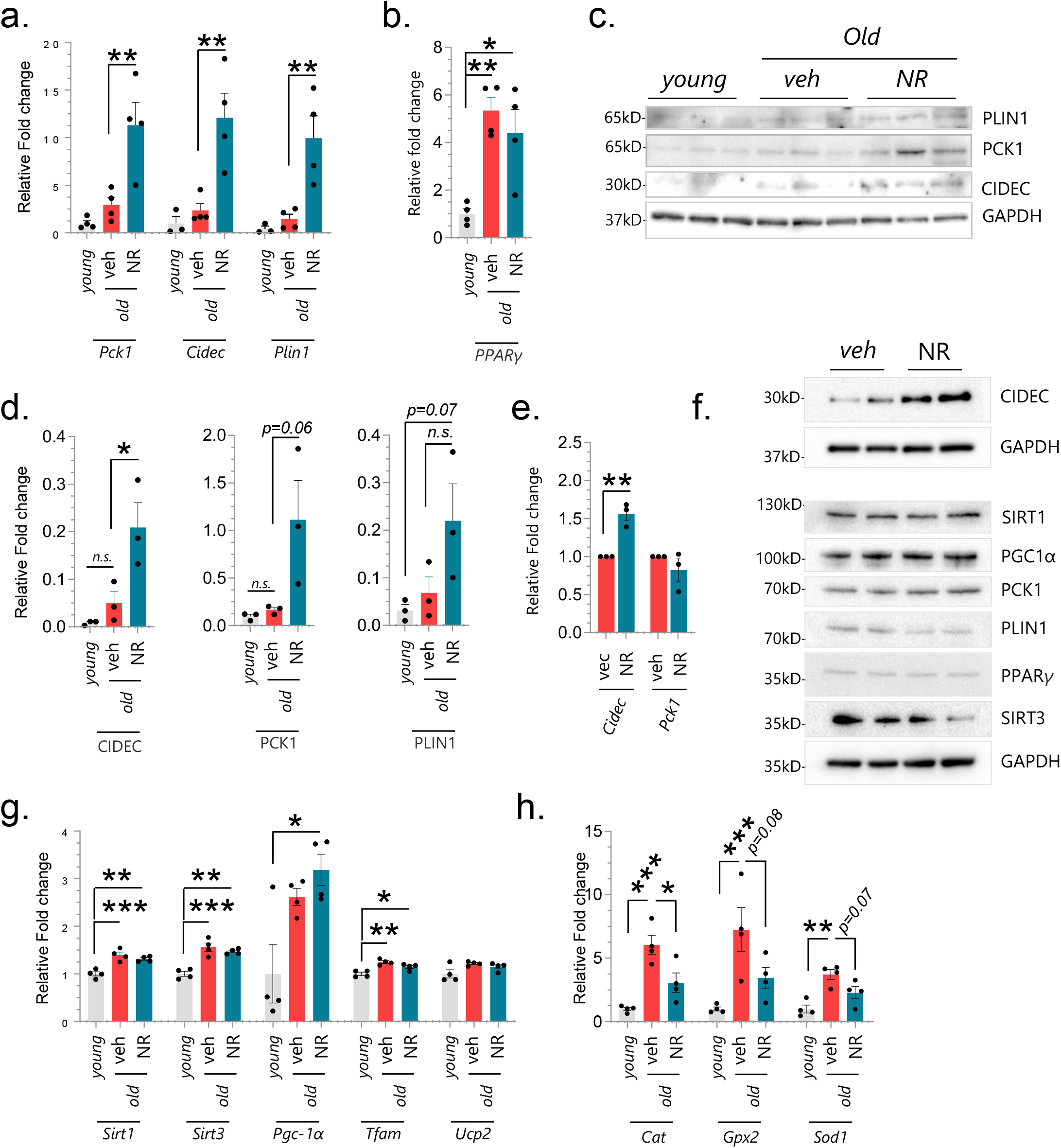
NAD+ supplementation using NR modulates the expression of CIDEC and PLIN1, key proteins of lipid droplets dynamics. **a-b**) Quantitative RT-PCR results demonstrate the relative fold change in Pck1, Cidec, Plin1, and PPARγ in the cochlea of young (2-month-old), old (12-month-old), and NR-treated old mice (12-month-old). N=4 mice per group, one-way ANOVA with Tukey’s post hoc test was used for statistical analysis). **c**) Western blot depicts protein expression in the cochlea of young (2-month-old), old (12-month-old), and NR-treated old mice (12-month-old). **d**) The graph demonstrates the quantification of the average signal from the Western Blot in Fig. 5c using the NIH ImageJ program. One-way ANOVA with Tukey’s post hoc test was used for statistical analysis. **e**) Quantitative RT-PCR results demonstrate the relative fold change in Pck1 and Cidec following NR treatment (1mM, 24H) in HEI-OC1 cells. **f**) Western blot depicts protein expression from HEI-OC1 cells following NR treatment (1mM, 24H) from two independent biological repeats (Lane 1 and 3 demonstrate experiment #1, Lane 2 and 4 demonstrate Experiment #2). **g-h**) Quantitative RT-PCR results demonstrate the relative fold change in genes in the cochlea of young (2-month-old), old (12-month-old), and NR-treated old mice (12-month-old). (N=4 mice per group, one-way ANOVA with Tukey’s post hoc test was used for statistical analysis). mean ± S.E., \**p* ≤ 0.05, \*\**p* ≤ 0.01, \*\*\**p* ≤ 0.001, and *n.s*., not significant.

Given the role of NAD+ in mitochondrial homeostasis, we also examined the expression levels of mitochondria-related genes (*Sirt1, Sirt3, Pgc-1α, Tfam, Ucp2*) in the cochlea and found an age-related elevation in their levels, with the exception of *Ucp2*. When treated and non-treated old cochlea were compared, no significant impact of NR was observed on mitochondria-related genes, which was in accord with RNAseq results in which no mitochondrial terms were identified in the top GO-term list from the cochlear samples (**Fig. 5g**). We observed similar results in cultured cochlear cells (**Fig. 5f** and **Supp. Fig. 3**).

The mtKeima strain was designed for ex-vivo imaging of mitochondria and mitophagy events *(16)*. It expresses a pH-dependent mtKeima protein and is resistant to lysosomal proteases. When mitochondria are in neutral pH, they fluoresce green but when in lysosomes, as during mitophagy, they are red. Thus, this strain permits the analysis of mitochondrial pools in and out of lysosomes. In our ex-vivo studies on live cochlear tissue, we show that NR treatment did not have a significant impact on mitophagy in auditory neurons (**Supp. Fig 4**). These results suggest that NR improves cochlear function with limited direct impact on mitochondria. However, lipid droplets are known to interact with mitochondria and reduce excessive mitochondrial reactive oxygen species overflow and fatty acid oxidation, contributing to the antioxidant capacity of cells *(25–27)*. Indeed, we observed that NR leads to a significant reduction in catalase (*Cat*), while there was a trend for a reduction in the levels of oxidative stress-related genes glutathione peroxidase 1 and superoxide dismutatase 1 (*Gpx*, and *Sod1*) that were elevated in the cochlea of old mice (**Fig. 5h**), suggesting an indirect effect of NR on mitochondrial homeostasis and function. Taken together, our results suggest a novel pathway of NR acting along the PPARγ-CIDEC, - PLIN1, -PCK1 axis, and that it potentially contributes to lipid droplet formation and protection of the cochlear cells from cytotoxic damage.

### NR halts the progression of hearing loss

We next tested if NR treatment was still beneficial for hearing loss after a hearing deficit had already developed. To address this, we first confirmed the substantial hearing loss in mtKeima mice at 15 m of age and then administered NR for 1.5 m (**Fig. 6a-b**). We found that NR had no significant impact on hearing thresholds at any given frequency (**Fig. 6b**), possibly due to a slight increase in ABR thresholds limiting the room for improvement. However, when individual threshold shifts were investigated, we found that NR treatment reduced threshold shifts significantly at 24 kHz, which only impacted females (**Fig. 6c** and **Supp. Fig. 5a**). These results show that NR treatment at a late stage may still be beneficial, at certain frequencies, even after hearing loss has already formed.

**Figure 6.**
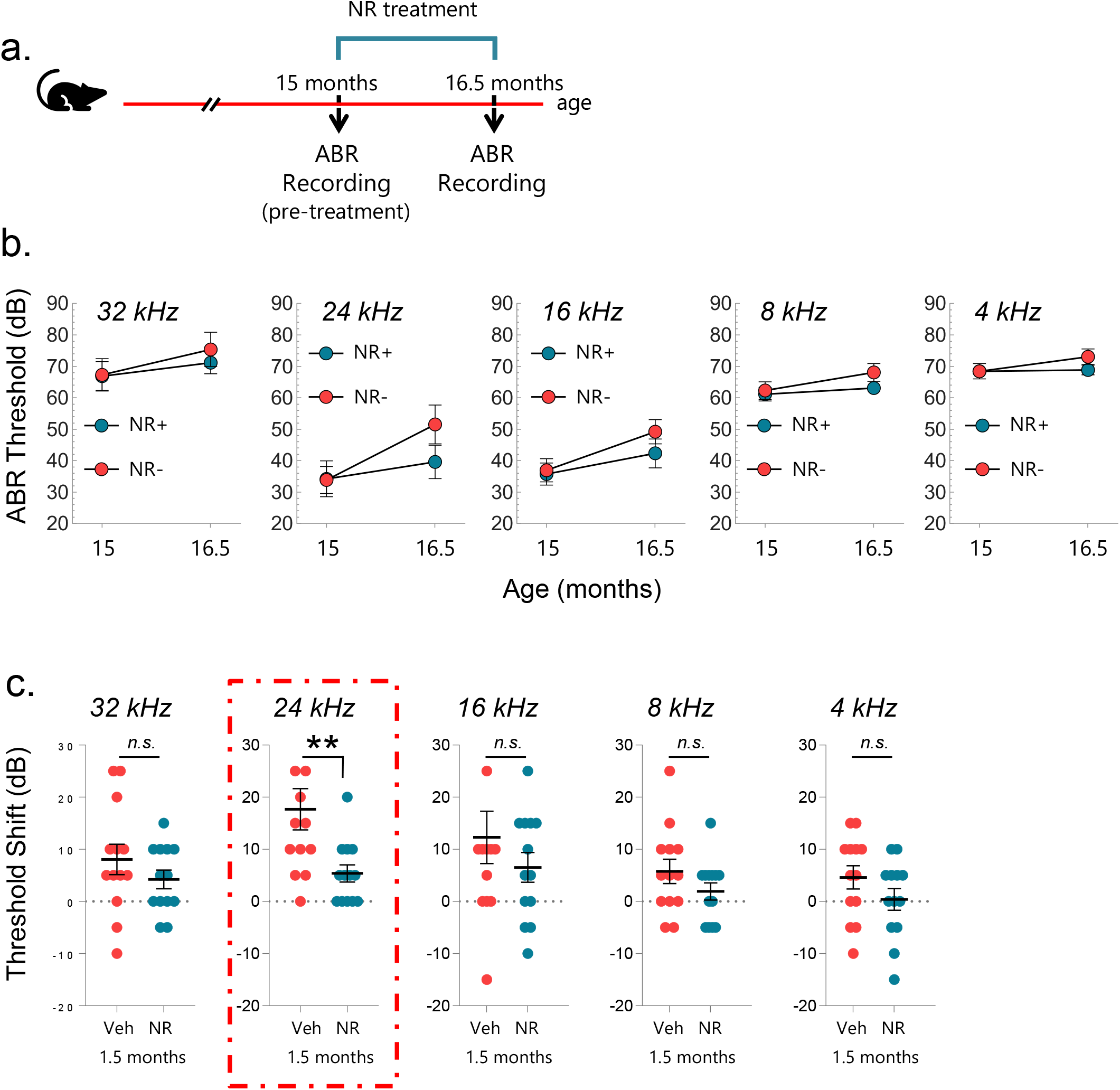
NAD+ supplementation using NR halts the further progression of ARHL in mice (mtKeima). **a**) Outline for NR treatment and ABR recordings in mice. **b**) ABR thresholds for NR-treated and non-treated WT mice at 15, and 16.5 m of age. A total of 26 WT mice were tested for hearing capacity at the age of 15 m and then split into two groups of NR-treated (N=13) and non-treated (N=13). Groups are gender-matched and hearing capacity-matched at 32 kHz at the age of 15 m. NR treatment started at the age of 15 m. ABRs in both groups were measured again at the age of 16.5 m, which corresponds to 1.5 m of NR treatment. Mixed effect analysis with Sidak’s multiple comparison test was used to determine significant differences. **c**) NR treatment prevents the increased threshold shift at 24 kHz. Note: ABR data in (b) were used to calculate the hearing shift. Two-tailed t-tests were used to determine significant differences. mean ± S.E., \**p* ≤ 0.05, \*\**p* ≤ 0.01, and *n.s*., not significant.

### Long-term NR administration prevents the progression of ARHL in WT mice with natural hearing loss

Our results so far show that NR administration prevents and halts the progression of ARHL in mice. To confirm these results, we also tested the impact of NR administration on ARHL using a different WT mouse strain (CBA/CaJ) that displays slower hearing loss progression than the mtKeima mouse strain and therefore better represents ARHL. Indeed, this model is widely used in auditory research, particularly for ARHL-related studies *(28)*. In accord with our previous results, we found that 24 m of NR administration prevented ARHL progression of high-frequency sounds (**Fig. 7a-b**). NR administration impacted hearing loss in female mice more prominently, although both sexes benefited (**Fig. 7b** and **Supp. Fig. 6 and 7)**. When hearing threshold shifts were compared, we found the NR-treated group developed less hearing loss per mouse compared to the non-treated group at 24 and 32 kHz (**Fig. 7c**). These results also demonstrate NR’s benefit on age-related hearing loss is not strain specific. Given the role of NR in synaptic connectivity (**Fig. 4c**), we next investigated NR’s effect on the integrity of synaptic transmission between sensory inner hair cells and auditory neurons. To address this, we analyzed synaptic ribbon counts in inner hair cells, which are electron-dense structures that tether synaptic vesicles at the presynaptic active zones and facilitate continuous synaptic transmission *(17, 29, 30)*. The cochlea consists of base, middle and apex regions. High-frequency sound is sensed at the base of the cochlea whereas low frequencies are sensed at the apex. We observed an age-associated decline in synaptic ribbon counts in the base and middle regions of the cochlea (**Fig. 8a-b**), consistent with the decline in ABR levels at high-frequency tones in the old mice (**Fig. 7b**). Remarkably, we found that NR restored the reduction in ribbon counts in the middle region of the cochlea, potentially contributing to hearing in the mid-frequency range (16-32 kHz) (**Fig. 8a**). However, no significant improvement was observed in the number of ribbon counts in the base region (**Fig. 8b**). This was rather surprising given that NR improves hearing at 32 kHz (**Fig. 7b**). One potential mechanism could be that NR exerts its impact in this region by preserving hair cell innervation by afferent auditory neurons, leading to improved synaptic transmission. Indeed, when orphan ribbons were compared (red only puncta in Figure 8a, b and d), we observed an age-related increase in the number of orphan ribbons (by 25% per inner hair cell) in the base region, which was fully reversed by NR treatment (**Fig. 8c**). The apex region, on the other hand, displayed no significant change in synaptic ribbon counts upon aging or with NR treatment (**Fig. 8d**).

**Figure 7.**
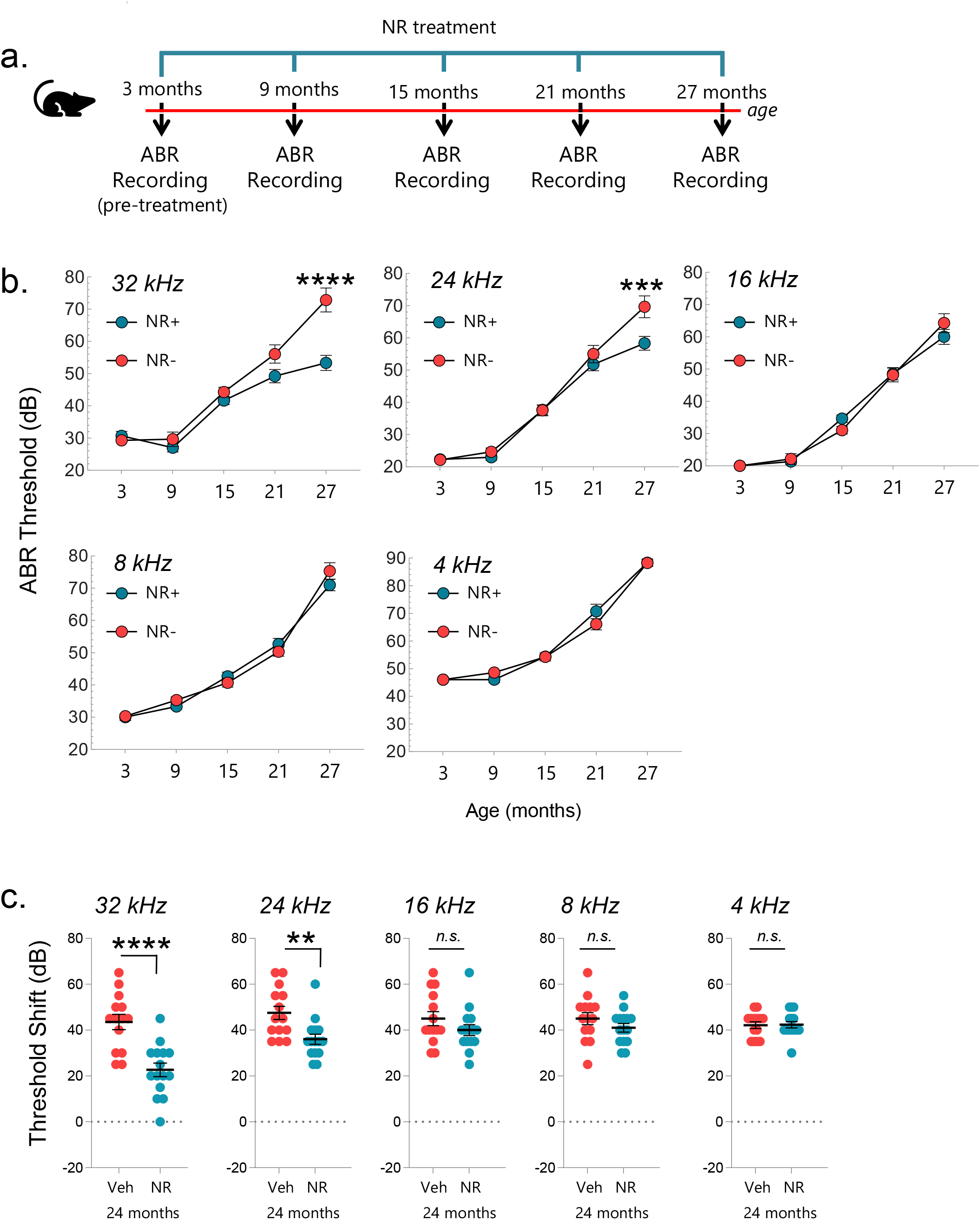
NAD+ supplementation using NR prevents the progression of age-related hearing loss in mice (CBA/CaJ). a) Outline for NR treatment and ABR recordings in mice. b) ABR thresholds for WT and NR-treated WT mice at 3, 9, 15, 21, and 27 m of age. A total of 29 WT mice were tested at the age of 3 m and then randomly split into two groups of NR-treated (N=15) and non-treated (N=14). NR treatment started at the age of 3 m. ABRs in both groups were measured again at the age of 9, 15, 21, and 27 m, which correspond to 6, 12, 18, and 24 m of NR treatment respectively. Mixed effect analysis with Sidak’s multiple comparison test was used to determine significant differences. c) NR treatment prevents the increased threshold shift at 24 and 32 kHz. Note: ABR data in (b) were used to calculate the hearing shift. Two-tailed t-tests were used to determine significant differences. mean ± S.E., \**p* ≤ 0.05, \*\**p* ≤ 0.01, \*\*\**p* ≤ 0.001, \*\*\*\**p* ≤ 0.0001 and *n.s*., not significant.

**Figure 8.**
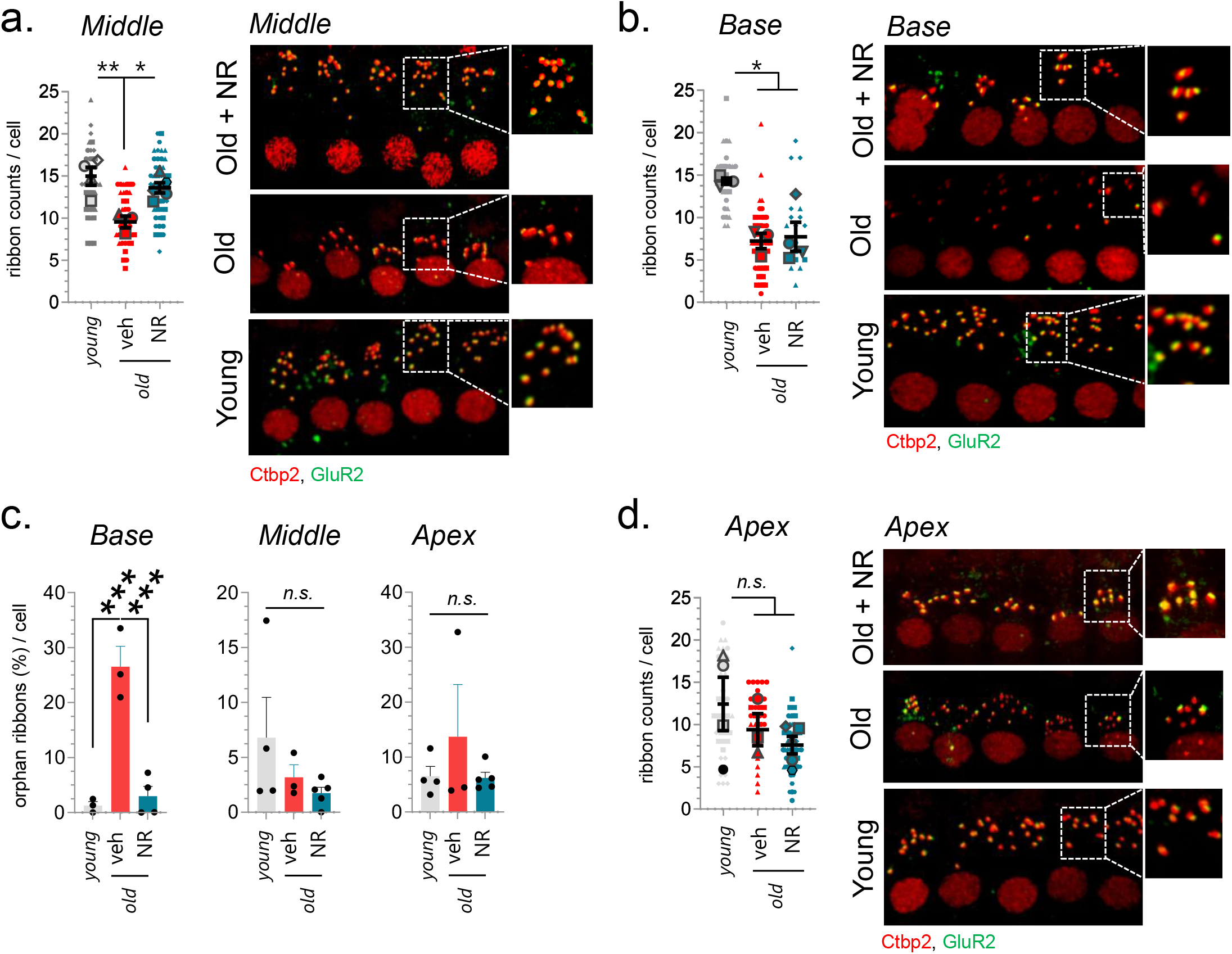
NAD+ supplementation rescues age-related decline in synaptic ribbon formation per inner hair cell in the cochlear middle region. The average synaptic ribbon count per inner hair cell in the cochlea middle region (a), base region (b), and apex region (d) is demonstrated on the left panels. The right panel shows a representative image of immunostaining for synaptic ribbons (red, anti-Ctbp2 (Ribeye) and post-synaptic receptor (green, anti-GluR2a) of cochlear middle segments (a), base segments (b), and apex segments (d). The anti-Ctbp2 faintly stains hair cell nuclei. Zoom-in images on the right-top corners of the panels illustrate the juxtaposition of ribbon–receptor pairs in selected areas. c) The graphs show the average number of orphan ribbons per inner cell. The red puncta (anti-Ctbp2) without green puncta (anti-GluR2a) juxtaposition is considered an orphan ribbon. mean ± S.E., one-way ANOVA with Tukey’s post hoc test was used for statistical analysis, mean ± S.E., \**p* ≤ 0.05, \*\**p* ≤ 0.01, \*\*\**p* ≤ 0.001, and *n.s*., not significant.

## DISCUSSION

ARHL is the most common type of sensorineural hearing loss. Current treatment approaches for ARHL have failed, including those targeting reactive oxygen species reduction. The slow progression of ARHL adds to the challenge for intervention because it requires treatment approaches safe for long-term use. Our previous findings showed that NAD+ supplementation using NR effectively prevented the progression of hearing loss in premature aging models *(8)*. NR can be administered orally and has no known serious side effects, and many safety studies have been done *(12)*, making it a good candidate for long-term administration. Most notably, using two different WT mouse strains, we found that long-term NR administration, for the duration of one or two years, partially prevents ARHL progression (**Fig. 1 and 7**). Our studies also show that NR administration halts further deterioration of already existing hearing loss at high frequencies (**Fig. 6**), providing a potential hearing loss treatment strategy for elderly individuals with some level of hearing loss.

We investigated potential underlying biological mechanisms of NAD+ repletion. To address this, we first performed cochlear transcriptomic analysis and found that several terms associated with synaptic transmission are up-regulated by NR treatment (**Fig. 4c**). In accordance with this, our in-depth wave-form analysis showed that NR prevents age-associated reduction in Wave I amplitude, which is indicative of the number of neurons firing between sensory and auditory nerve cells (**Fig. 3**). Indeed, we observed that NR treatment rescues the age-related decline in synaptic ribbon counts in the middle turn of the cochlea (**Fig. 8a**). Synaptic ribbons are largely composed of RIBEYE (Ctbp2) proteins through multiple RIBEYE-RIBEYE interactions *(31)*. Interestingly, RIBEYE has NAD+/NADH binding pocket that modulates synaptic ribbon assembly and activity, suggesting a potential mechanism of action of NR. However, although NR improved high-frequency hearing (**Fig. 7b**), it did not increase the synaptic ribbon counts in the base region that corresponds to sensing high-frequency tones (>=32 kHz) (**Fig. 8b**). Therefore, we speculated that NR benefits hearing through additional mechanisms besides upregulation of ribbon counts. Indeed, a previous study showed that NR prevents noise-induced neurite retraction from inner hair cells *(7)*. In line with these findings, we found that NR prevented an age-related increase in the number of orphan ribbons specifically in the base region, suggesting that NR preserves the innervation of inner hair cells in this region. We propose that the increased number of orphan ribbons with age reflects a deterioration of afferent auditory neuron dendrites, and that the reduction in orphan ribbons (and correspondingly improved Wave I amplitudes) with NR treatment reflects preservation of afferent auditory neuron innervation and function. Given the decrease in orphan ribbons, but no change to hair cell ribbon counts in the base, we speculate that the main target of NR in this region is the auditory afferent neurons. Altogether, our results suggest that NR benefits hearing by contributing to synaptic stability between sensory cells and auditory primary neurons. However, we do not rule out the possibility that NR impacted other regions along the auditory pathway. In fact, we observed that NR also significantly improved Wave-III amplitude, which represents the number of neurons firing in a superior olivary complex in the brainstem.

Besides the NR-mediated upregulation of synaptic transmission, our transcriptomic analysis also indicates molecular signatures of lipid droplet regulation as a potential underlying mechanism of NR’s benefit on hearing loss. Lipid droplets have long been considered lipid storage units of the cells, but it is now more apparent that their expansion and shrinkage are highly dynamic and tightly coupled to their interaction with other organelles to control energy homeostasis and cellular stress. For instance, lipid droplets accumulate free acids in their units and move them into mitochondria when needed for energy demand, reducing free acid in the cytosol and hence lipotoxicity while contributing to energy homeostasis *(32)*. Here, we show for the first time that NAD+ supplementation using NR elevates the expression of key proteins of lipid droplet dynamics, CIDEC and PLIN1, in the cochlea. CIDEC protein binds to lipid droplets and regulates their enlargement *(22)*. PLIN1, a lipid droplet coating protein, binds to CIDEC and regulates lipid storage and lipolysis *(22, 33)*. PLIN1 directly interacts with Mfn2, a mitochondrial fusion protein, and facilitates the contact between mitochondria and lipid droplets, mediating lipolytic processes and cellular metabolism *(34)*. In accord with this, recent data showed that patients with Mfn2 variant (D414V) exhibit a hearing loss phenotype *(35)*. NR-mediated increases in CIDEC and PLIN1 hence suggest a potential regulation of lipid droplet dynamics by NAD+. However, the molecular basis of how NR modulates CIDEC and PLIN1 is not yet clear. *Cidec* and *Plin1* gene promoters contain functional PPAR-responsive elements and are targeted by PPARγ *(36)*. PPAR activation protects the cochlea from oxidative stress *(19)*. We identified the PPAR signaling pathway in NR-treated cochlea using RNA seq analysis (**Fig. 4d**). Interestingly, it was shown that the inhibition of NAD+ synthesis lowers α-ketoglutarate-mediated PPAR expression in 3T3-L1 cells *(37)*. However, we did not observe a significant change in the levels of PPARγ in NR-treated cochlea or cochlear cells, suggesting that NR might modulate PPARγ activity or act downstream of the PPARγ pathway. These observations warrant further investigations to elucidate the effects of NR on lipid droplet dynamics and its impact on hearing loss.

Our results show that NR particularly prevents hearing loss at higher frequencies while its impact fades with decreasing frequency, suggesting that NR predominantly modulates the cochlear middle and base segments. These results may potentially reflect higher firing rates for basal hair cell synapses, and therefore higher metabolic demand and excitotoxicity for higher frequency neurons *(38)*. Alternatively, NR might not adequately reach the apex region. In a different study, IP-injected NR exerted its benefits on the apex region after noise exposure *(39)*. Thus, we believe that NR reaches all regions of the cochlea although the administered NR dosage might not be adequate to exert its benefits in the whole cochlea. Alternatively, NR might regulate biological pathways predominately in the base area of the cochlea. For instance, SOD2 protein expression shows a base-to-apex increasing gradient in afferent auditory neurons in rodent cochlea *(40)*. SOD2 is an antioxidant enzyme inside mitochondria whose levels increase with NR treatment *(41)*. Interestingly, mitochondria expressing more SOD2 show a closer association with lipid droplets *(42)*. Regardless, NR’s specific impact on high frequencies also suggests that NR’s benefit on hearing loss primarily originates from its effect on the cochlea. Taken together, our study demonstrates the therapeutic potential of NAD+ repletion, using NR, for the treatment of ARHL via improving the synaptic transmission in the cochlea and it points out lipid droplet dynamics as novel NR targets in the cochlea.

## METHODS

### Cell Culturing

HEI-OC1 cells were a generous gift of Dr. Federico Kalinec and their maintenance was previously described*(43)*. Briefly, we cultured HEI-OC1 cells in Dulbecco’s modified Eagle medium (DMEM) containing 10% fetal bovine serum (FBS) in a humidified chamber under permission conditions (10% CO_2_ at 33°C).

### Animals

mtKeima transgenic mice were a generous gift of Dr. Toren Finkel and described previously *(16)*. These mice (http://www.informatics.jax.org/allele/MGI:5660493) are on a mixed genetic background and to rule out the cadherin23 AHL allele as a potential confounder in our analyses, mice tested and found to be all WT for the cadherin23 gene. Additionally, mice were genotyped for the C57BL/6J nicotinamide nucleotide transhydrogenase gene (NNT) mutation and were WT.

NR was given orally in drinking water to mouse strains at a concentration of 7 mg/ml (24 mM), while the non-treated control groups received regular drinking water. The justification for the NR concentration used in animal studies was described previously *(8)*. The water bottles, with or without NR, were changed twice per week. 12-month-old mtKeima mice were used for quantifying NAD^+^ levels in the cochlea. NR administration to mtKeima mice started at 2 m and treatment lasted for 10 m until mice were 12 m of age; or started at the age of 15 m and lasted for 1.5 m until 16.5 m of age. The cochlea were dissected for further assessment at the end of the NR administration right after the auditory assessment. The WT CBA/CaJ mice were purchased from The Jackson Laboratory (cat number #000654). NR administration to WT CBA/CaJ mice started at 3 m of age and lasted for 24 m until mice were 27 m of age. Mice were maintained on a 12h light-dark cycle and fed ad libitum. All animal protocols were approved by the Animal Care and Use Committee of the Intramural Research Program of the National Institute on Aging, OSD-361-2023 in accordance with the National Research Council’s Guide for the Care and Use of Laboratory Animals.

### NAD+ quantification

Dissected cochlea were placed in NADH/NAD extraction buffer (Abcam, ab65348) and homogenized with a micro pestle. NAD+ and NADH levels in cochlea were quantified using the NAD/NADH Assay Kit per manufacturer instructions. Samples were normalized to total protein concentration in each cochlea using Pierce™ BCA Protein Assay Kit.

### Audiometry

ABR methodology has been described previously*(8)*. Briefly, mice were anesthetized with ketamine (100 mg/kg) and xylazine (10 mg/kg) via an intraperitoneal (i.p.) injection and placed in a soundproof chamber on a heating pad in such a way that the recorded ear was 7 cm away from the sound source (MF1 Multi-Field Magnetic Speaker). After inserting the needle electrodes sub-dermally (vertex–ventrolateral to pinna), tone burst stimuli (5 ms duration with a 0.1-ms rise-fall time) were presented at variable volume (10–90 dB SPL) in 5 dB steps at 4, 8, 16, and 32 kHz using RZ6 system (Tucker Davis Technologies) with Biosig software (Tucker Davis Technologies). The minimum volume threshold (in dB) that evokes a response at a given frequency was recorded as the outcome measure. The waveforms were determined by an average of 512 responses. The ABR threshold was determined by visual inspection and considered to be the lowest stimulus level at which at least one wave was present.

DPOAE measurement was described previously *(8)*. Briefly, an earplug connected to a small microphone (ER-10B+) and two speakers (MF1 Multi-Field Magnetic Speaker) was inserted into the outer ear canal of each mouse. Using the RZ6 system (Tucker Davis Technologies) with Biosig software (Tucker Davis Technologies), a series of auditory stimuli were delivered to the speaker, each composed of two tones at equal decibel levels but distinct frequencies, *f*1 and *f*2, where *f*2 > *f*1, *f*2/*f*1 = 1.2 at *f*0 = 10, 12, 16, and 32 kHz (*f*0 = (*f*1 × *f*2)1/2). The decibel level of both tones varied over the range of 80 dB SPL to 10 dB SPL in 5-dB steps. The distortion product at the frequency 2*f*1 − *f*2 was recorded at each frequency tested as an average of 512 responses.

### ABR Waveform Reconstruction

Raw ABR recording data were extracted using the BioSigRZ software. Voltage values were sampled at a rate of 50,000 Hz (every 0.02 ms) for a duration of 4.5 ms following stimulus presentation. Representative waveforms were calculated and reconstructed offline by averaging the voltage values at each time point using TDT BioSigRZ software. Waves I-V were identified by a series of characteristic peak-to-following-trough forms.

### RNA Isolation and Quantitative real-time PCR (qPCR)

Total RNA from cochlear tissue was extracted using Trizol Reagent (Zymo Research, #R2050-1-50) and Direct-zol RNA Miniprep (Zymo Research, #R2050) as described previously*(44)*. Next, one microgram of isolated RNA was reverse transcribed using the iScript™ cDNA Synthesis Kit (BioRad). Using the DyNAmo HS SYBR green qPCR kit (F-410L, ThermoFisher Scientific) with the CFX connect real-time PCR detection system (Bio-Rad), qPCR was performed. Primer sequences are listed in Supplementary Table S1. Experimental values were normalized to values for GAPDH. The same protocol was followed for RNA isolation from cochlear cells except those steps to break the cochlear bone were skipped.

### RNA sequencing

RNA from mtKeima mice cochlea was isolated as described above. Library construction and sequencing were performed by Novogene. RNA purity was checked using the NanoPhotometer® spectrophotometer (IMPLEN, CA, USA). RNA integrity and quantitation were assessed using the RNA Nano 6000 Assay Kit of the Bioanalyzer 2100 system (Agilent Technologies, CA, USA). A total amount of 1 μg RNA per sample was used as input material for the RNA sample preparations. Sequencing libraries were generated using NEBNext® UltraTM RNA Library Prep Kit for Illumina® (NEB, USA) following the manufacturer’s recommendations and index codes were added to attribute sequences to each sample. Briefly, mRNA was purified from total RNA using poly-T oligo-attached magnetic beads. Fragmentation was carried out using divalent cations under elevated temperature in NEBNext First Strand Synthesis Reaction Buffer (5X). First-strand cDNA was synthesized using random hexamer primers and M-MuLV Reverse Transcriptase (RNase H-). Second strand cDNA synthesis was subsequently performed using DNA Polymerase I and RNase H. Remaining overhangs were converted into blunt ends via exonuclease/polymerase activities. After adenylation of 3’ ends of DNA fragments, the NEBNext adaptor with a hairpin loop structure was ligated to prepare for hybridization. To select cDNA fragments preferentially of 150∼200 bp in length, the library fragments were purified with the AMPure XP system (Beckman Coulter, Beverly, USA). Then 3 μl USER™ Enzyme (NEB, USA) was used with size-selected, adaptor-ligated cDNA at 37 °C for 15 min followed by 5 min at 95 °C before PCR. Then PCR was performed with Phusion High-Fidelity DNA polymerase, Universal PCR primers, and Index (X) Primer. At last, PCR products were purified (AMPure XP system) and library quality was assessed on the Agilent Bioanalyzer 2100 system. The clustering of the index-coded samples was performed on an Illumina Novaseq sequencer according to the manufacturer’s instructions. After cluster generation, the libraries were sequenced on the same machine and paired-end reads were generated.

RNAseq analysis was provided by Novogene. Original image data file from Illumina was transformed to sequenced reads (raw data) by CASAVA base recognition (base calling). Datasets are being uploaded to GEO. Raw data was subjected to data QC including error rate distribution, GC content distribution, and data filtering. Mapping was performed using STAR (v2.6.1d, mismatch=2) and reads were mapped to the mouse reference genome mm10. Quantification was conducted by FeatureCounts (v1.5.0) under default mode. Normalization and differential gene expression used DESeq2 (v1.26.0) *(45)*. Genes with p-value ≤0.05 were passed on for enrichment analysis (Gene Ontology and KEGG) using ClusterProfiler (v3.8.1) and terms with padj <0.05 were considered significant.

### Reagents and Immunoblotting

Cochlear tissue and cells were lysed using RIPA buffer (Cell Signaling, #9806) supplemented with Halt™ Protease and Phosphatase Inhibitor Cocktail (Thermo Scientific™, #78444). Western blotting was performed according to the manufacturer’s instructions. Briefly, protein concentration was measured using a Pierce™ BCA protein assay kit (Thermo Fisher, 23225). Samples were separated on 4-12% Bis-Tris gel (Thermo Fisher Scientific, #NP0336BOX) and transferred to PVDF membranes (BioRad, #1620177). Unless indicated otherwise, membranes were blocked in Tris Buffered Saline (TBS) with 0.1% Tween 20 TBST + 3% milk at room temperature for 1H, incubated overnight with primary antibodies, washed 3X with TBST for 5 mins, incubated with HRP-conjugated secondary antibody, and developed using SuperSignal™ West Femto Maximum Sensitivity Substrate (Thermo Scientific™, 34095) according to manufacturer’s instructions. Antibodies were used according to the manufacturer’s instructions to detect the following antigens: CIDEC/Fsp27 (Ptglab, #12287-1-AP), PCK1 (Ptglab, #16754-1-AP), Plin1 (Abcam, Ab3526, 1:1000), Sirt1 (Santa Cruz, sc15404, 1:1000), GAPDH (Abclonal, AC027, 1:5000), PPARγ (Abcam, ab59256, 1:1000), CtBP2 IgG1 (BD Biosciences, 612044, 1:200), GluR2 IgG2a (Millipore, MAB397, 1:1000), myosin VIIa (Novus Biologicals, NB120-3481, 1:200).

### Cochlear immunofluorescence and imaging

Isolated cochlea were punched into oval and round windows using a syringe needle and rinsed/fixed with 4% paraformaldehyde in 1X cold PBS. Then, cochlea were further fixed in 4% paraformaldehyde in 1X cold PBS for additional 2 hours at 4C^0^. Following fixation, samples were rinsed 3 × 20 min in PBS and dissected under a stereomicroscope to the three turns: apical, middle and basal. Tissues were permeabilized in PBS + 0.25% Triton X-100 solution for 10 min at room temperature on a rocking platform; blocked with 10 % goat serum and 25 mM glycine for 1 h at RT. Tissues were incubated at 4°C overnight with the following primary antibodies: monoclonal mouse anti-carboxyl-terminal binding protein 2 (CtBP2) IgG1 at 1:200 (612044; BD Biosciences) counterstained with goat anti-Mouse IgG1 conjugated with Alexa Fluor 568 (#A-21124), monoclonal mouse anti-GluR2 IgG2a at 1:1000 (MAB397; Millipore) counterstained with goat anti-Mouse IgG2a conjugated with Alexa Fluor 488 (#A-21131), and polyclonal rabbit anti-myosin VIIa at 1:200 (NB120-3481; Novus Biologicals) counterstained with goat anti-Rabbit IgG (H+L)conjugated with Alexa Fluor 647 (#A-21244). Antibodies were added with 1% goat serum. The following day, after three 15-min PBS washes, the tissues were incubated with the Alexa-conjugated secondary antibodies at a concentration of 1:600 for 1 h in darkness at room temperature. Following the final washes after secondary incubations, samples were carefully mounted on slides using ProLong Glass antifade media and left to dry for at least 24 h before image acquisition. Frequency regions corresponding to 16 and 32 kHz were located through their distance from cochlear apex, based on the place-frequency map from Müller et al. *(46)* and imaged with a 63x 1.4NA Plan Apo objective on a Zeiss 880 LSM Airyscan confocal microscope with a 47nm xy pixel size (Carl Zeiss, Oberkochen,Germany). After acquisition, images were Airyscan processed.

### Ex-vivo mitophagy analysis in the cochlea

Ex-vivo mitophagy analysis in mtKeima mice has been described previously *(16, 47)*. Ex-vivo mitophagy analysis on cochlear tissues was performed as follows: Dissected temporal bone is placed in a silicone elastomer-coated dissection dish filled with 1X PBS at 4°C. Cochlea with otic capsule was removed from temporal bone and stabilized on the dish using pins. The otic capsule was slowly snipped off using Dumont #5 Fine Forceps (Fine Science Tools, #11254-20) and Vannas-Tübingen Spring Scissors (Fine Science Tools, #15003-08) and separated from cochlea inside the capsule without damaging the cochlear structure. The cochlea is then fragmented into the apex, middle, and base and placed on Nunc™ Glass Bottom Dishes (Thermo Scientific™, #150680) with 1X PBS at 4°C containing DAPI (Thermo Scientific™, #62248) at a final concentration of 1/3000 mg/ml. Following 10 mins of incubation, excess PBS was removed from the plate, and cochlea were imaged using a Zeiss 880 LSM confocal microscope.

### Statistical Analysis

Two-tailed *t*-tests were used to determine the differences between the two groups while One-way ANOVA with Tukey’s post hoc test was used to determine significant differences across multiple samples unless indicated otherwise in figure panels. Mixed effect analysis with Sidak’s multiple comparison test was used to determine significant differences in ABR thresholds in NR treated and non-treated groups. Statistical analyses were performed with GraphPad Prism version 7 (GraphPad Software, Inc.).

## ACKNOWLEDGEMENT

We would like to thank Dr. Federico Kalinec for generously providing HEI-OC1 cells. We also thank Dr. Nuo Sun and Dr. Toren Finkel for providing mtKeima transgenic mice. Lastly, we thank Dr. Xuili Dan for providing her expertise in mitophagy score analysis in the mtKeima mice cochlear tissues.

## Supplementary Tables and Figures

**Supplementary Figure S1.**
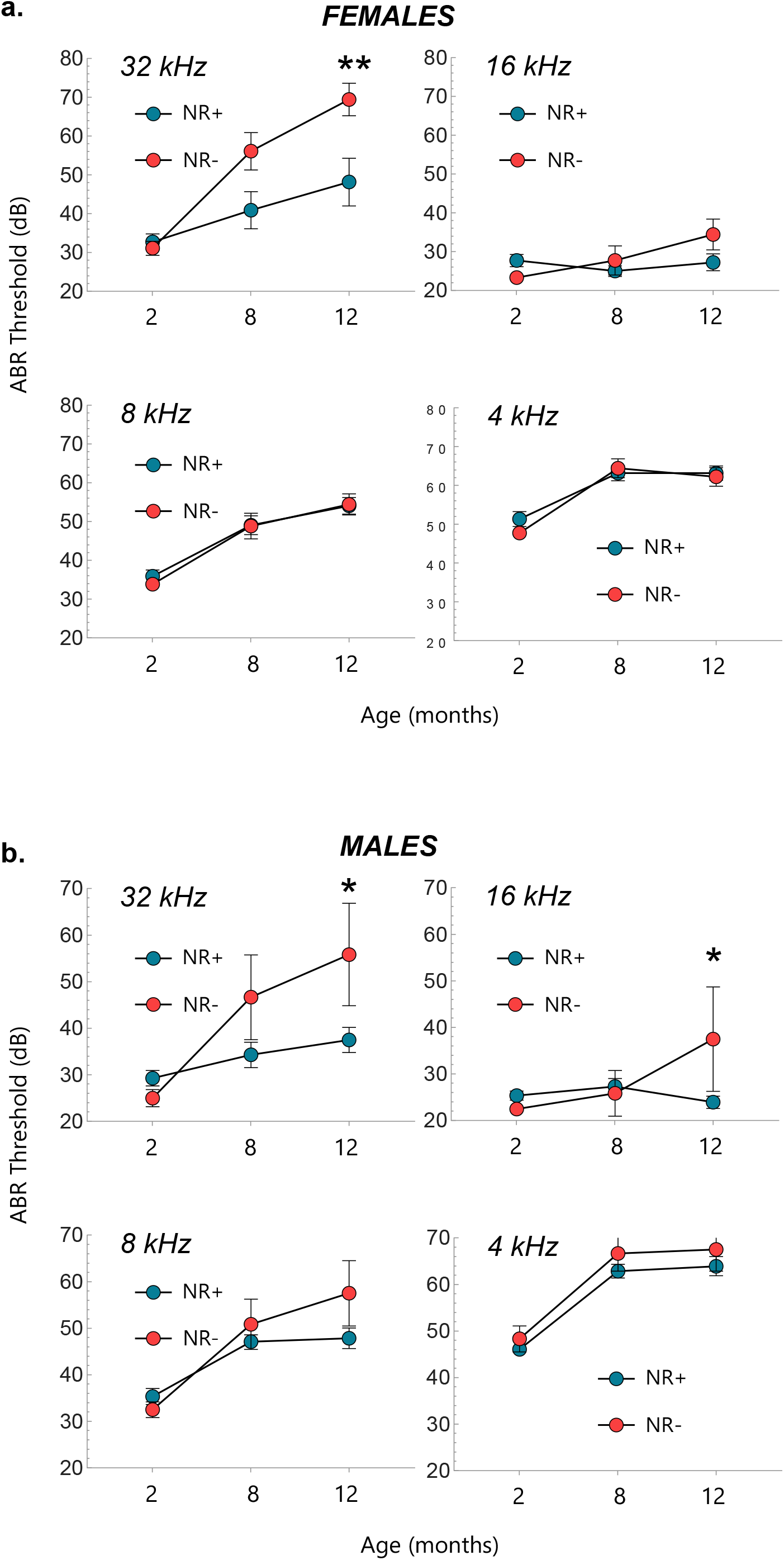
ABR thresholds at 4, 8, 16, and 32 kHz for NR-treated and non-treated female (a) and male (b) mice at 2, 8, and 12 m of age are shown. mean ± S.E., \**p* ≤ 0.05, \*\**p* ≤ 0.01, \*\*\**p* ≤ 0.001, \*\*\*\**p* ≤ 0.0001 and *n.s*., not significant.

**Supplementary Figure S2.**
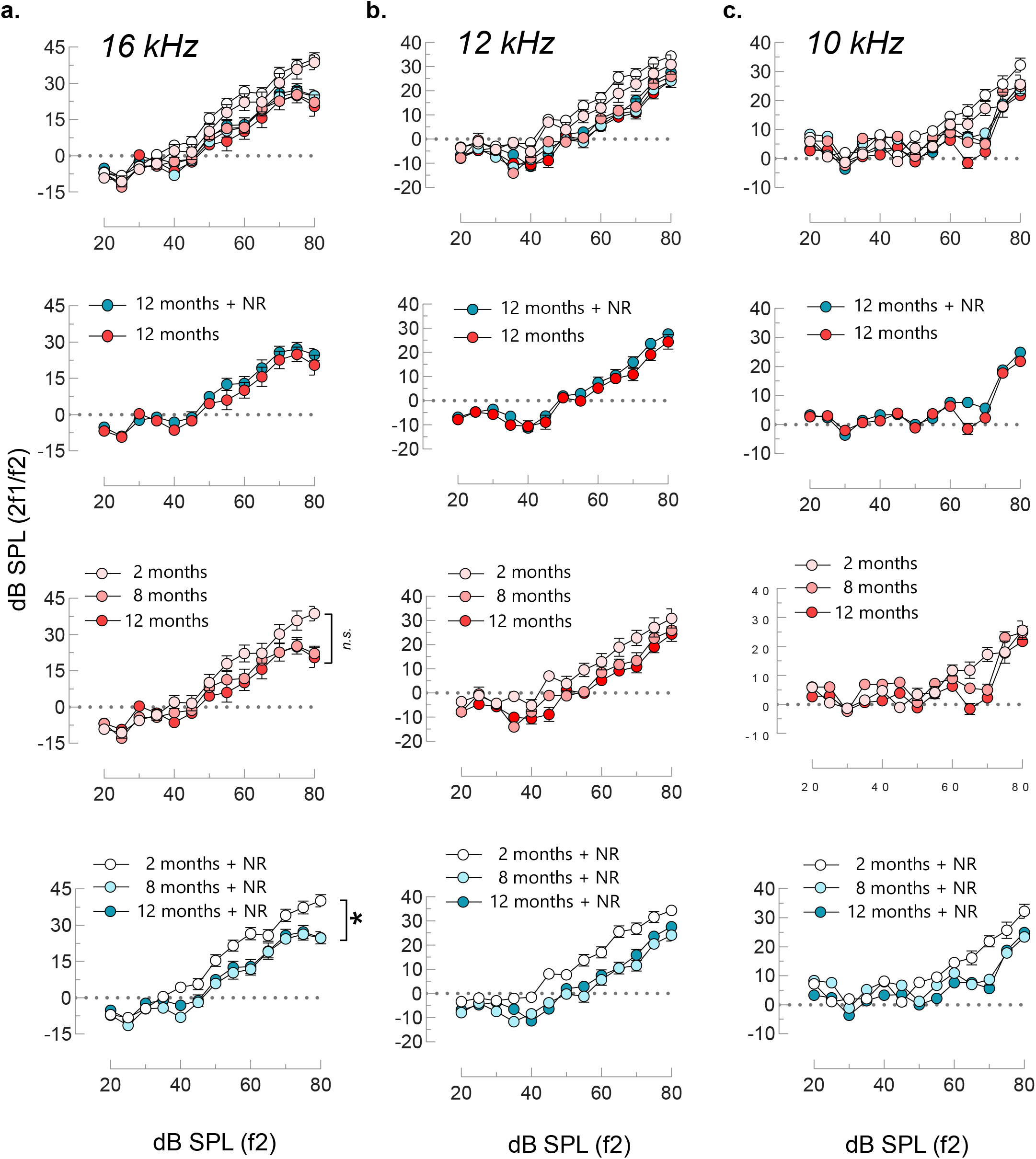
DPOAE levels at 16 kHz (a), 12 kHz (b) and 10 kHz (c) are shown at the age of 2, 8, and 12 m of age in NR-treated (N=25) and non-treated mice (N=13). The area under the curve (above −10 on the *y*-axis) is calculated for each sample and two-way ANOVA with Tukey’s post hoc test was used for statistical analysis. mean ± S.E., \**p* ≤ 0.05, \*\**p* ≤ 0.01, \*\*\**p* ≤ 0.001, \*\*\*\**p* ≤ 0.0001 and *n.s*., not significant.

**Supplementary Figure S3.**
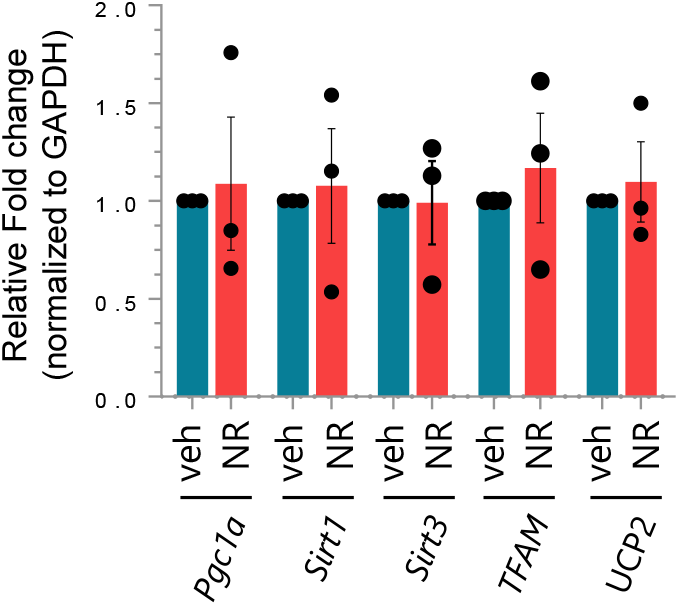
Quantitative RT-PCR results demonstrate the relative fold change in *Pgc1a, Sirt1, Sirt3, TFAM*, and *UCP2* following NR treatment (1mM, 24H) in HEI-OC1 cells. Two-tailed t-tests were used to determine significant differences. mean ± S.E.

**Supplementary Figure S4.**
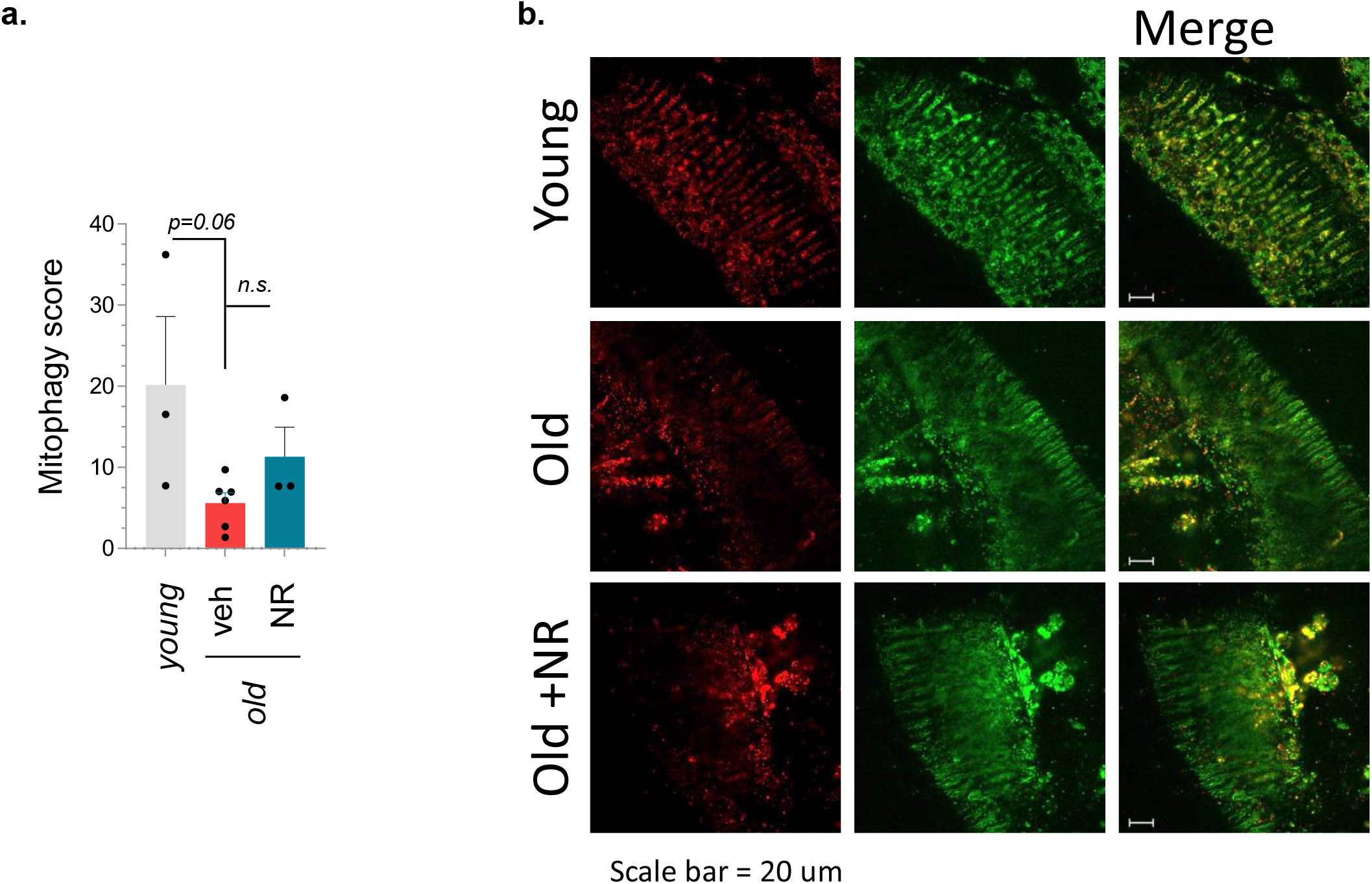
NR administration has no significant impact on *In-vivo* mitophagy score in cochlear tissues. The graph shows mitophagy scores in cochlear tissues freshly dissected from young (non-treated), old (non-treated), and old (NR-treated) mtKeima mice. The mitophagy score is calculated by quantifying the ratio of the lysosomal signal (red, 561 nm) to the mitochondrial signal (green, 458 nm) using Zeiss ZEN software. One-way ANOVA with Tukey’s post hoc test was used for statistical analysis. See the Methods section for details.

**Supplementary Figure S5.**
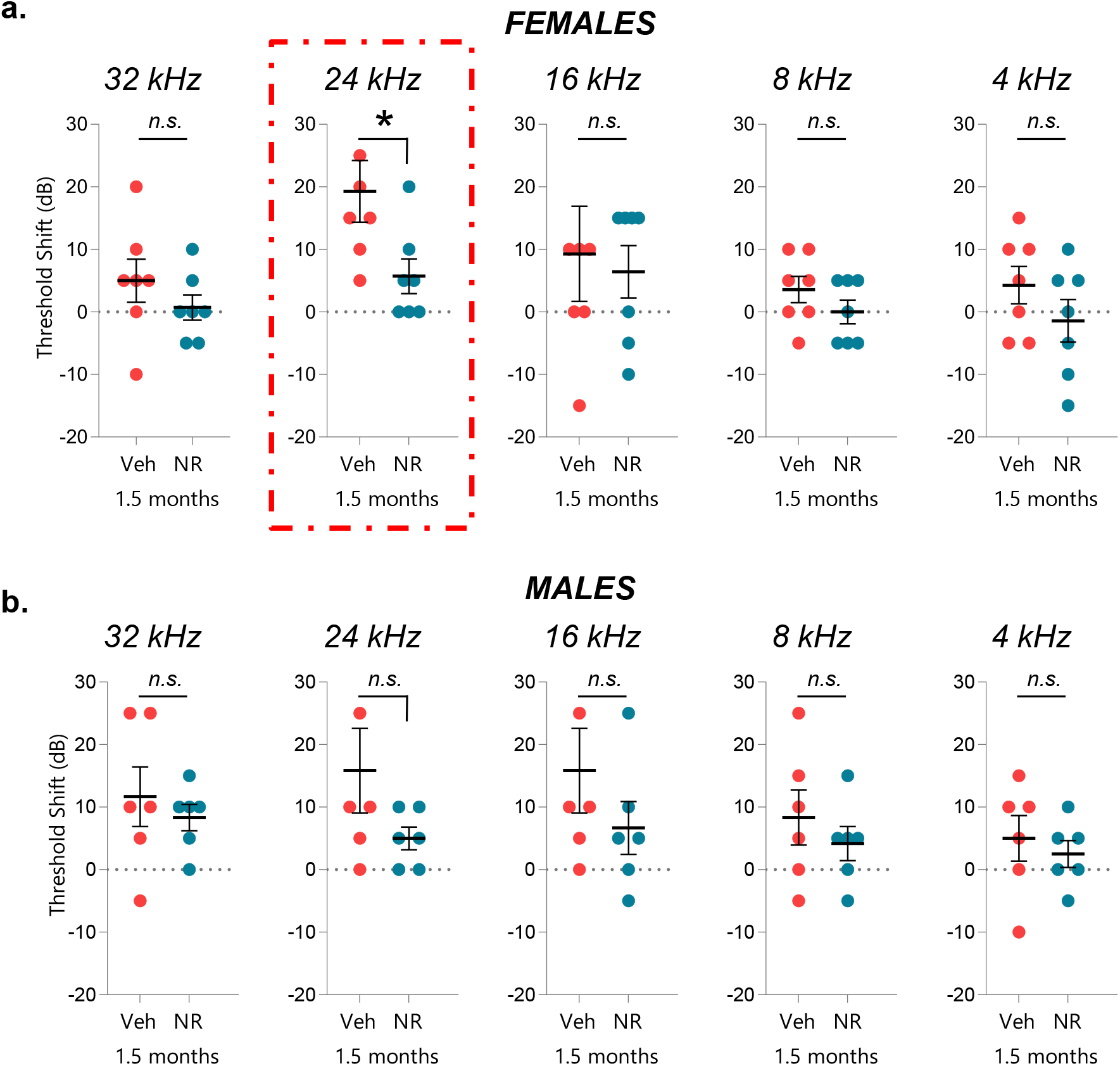
The graphs show the gender-specific demonstration of the threshold shifts in female (a) and male (b) mice in Figure 6. Two-tailed t-tests were used to determine significant differences. mean ± S.E., \**p* ≤ 0.05 and *n.s*., not significant.

**Supplementary Figure S6.**
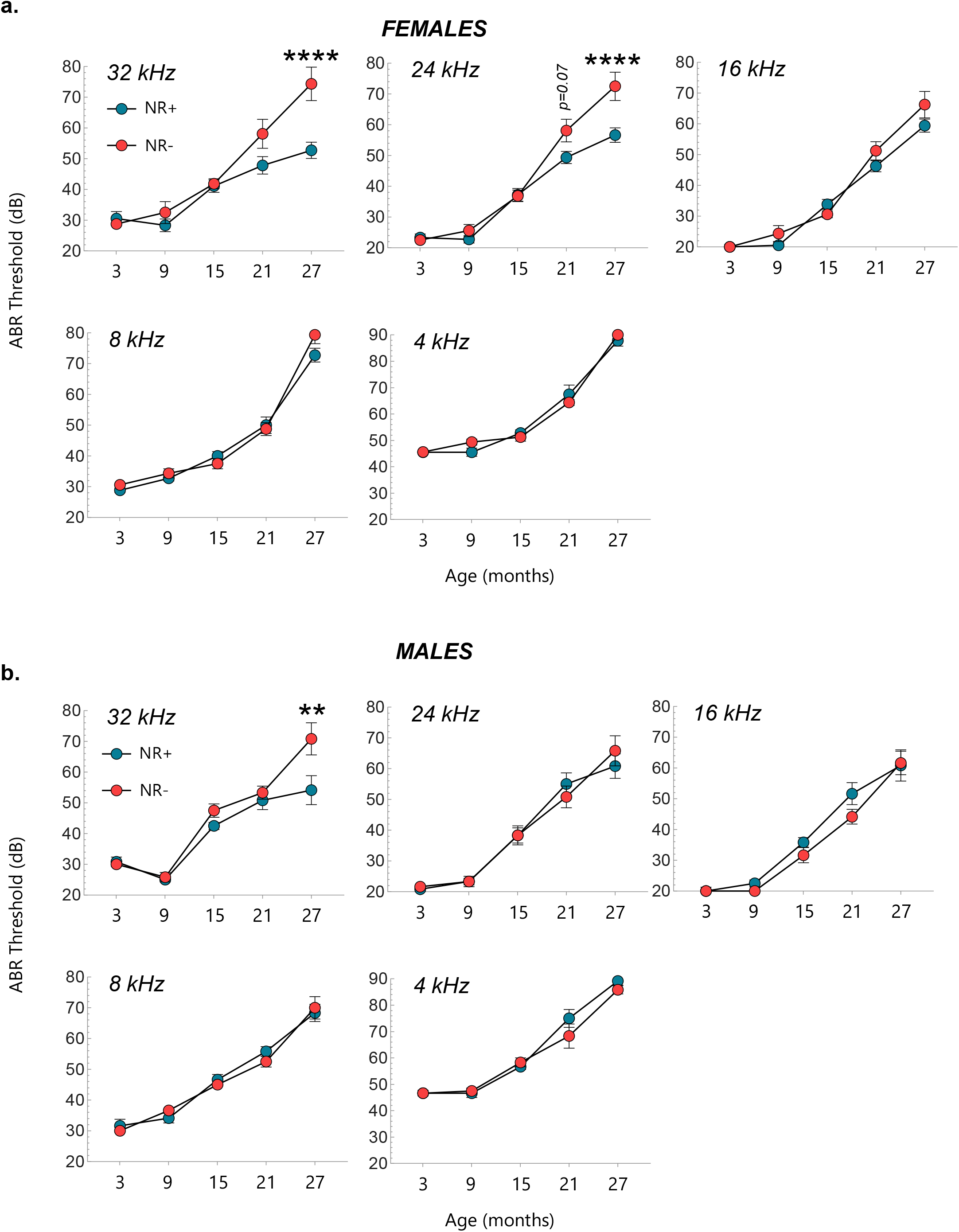
The graphs show the gender-specific demonstration of the ABR thresholds in female (a) and male (b) mice in Figure 7. Mixed effect analysis with Sidak’s multiple comparison test was used to determine significant differences. mean ± S.E., \**p* ≤ 0.05, \*\**p* ≤ 0.01, \*\*\**p* ≤ 0.001, \*\*\*\**p* ≤ 0.0001 and *n.s*., not significant.

**Supplementary Figure S7.**
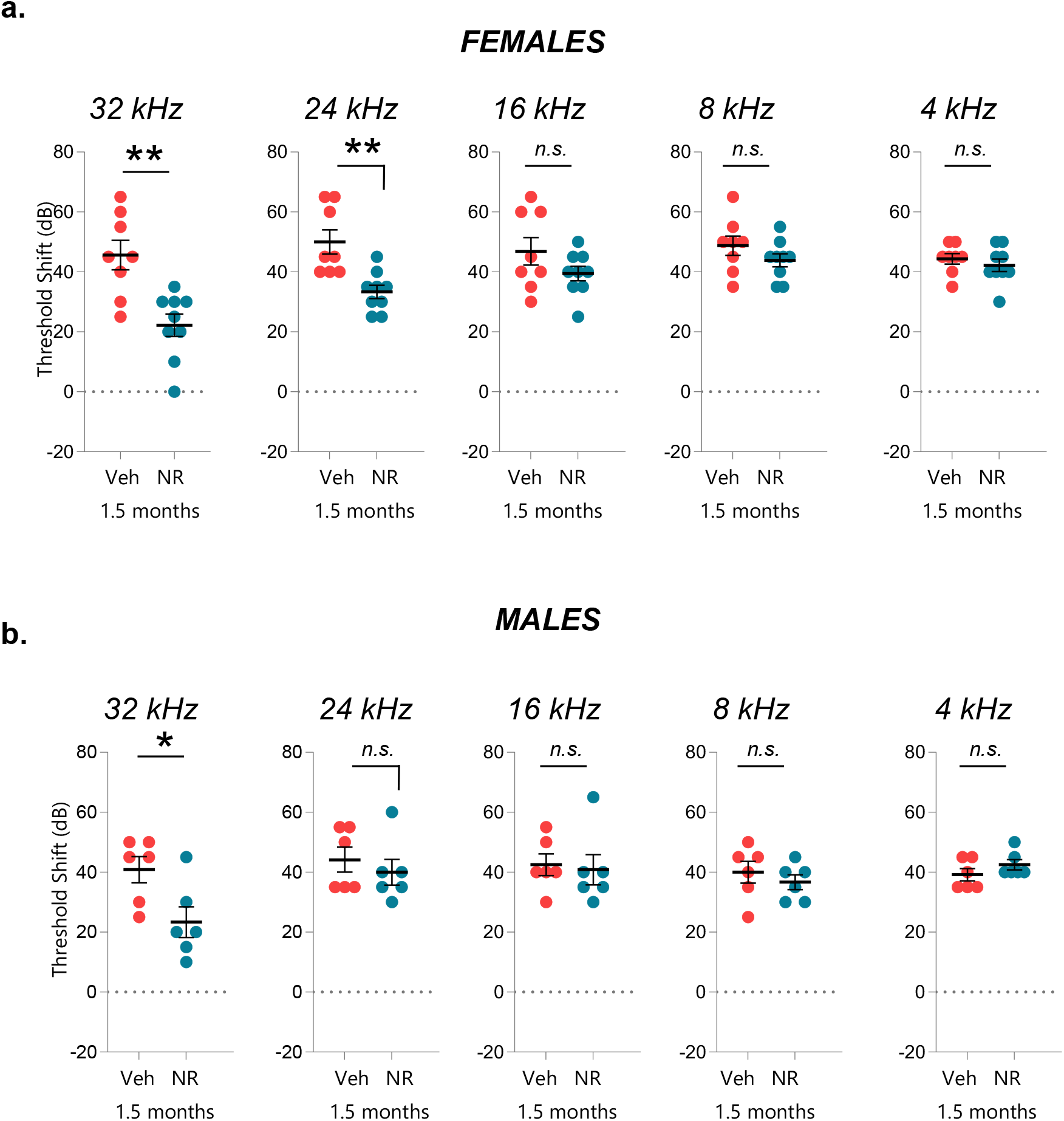
The graphs show the gender-specific demonstration of the threshold shifts in female (a) and male (b) mice in Figure 7. Two-tailed t-tests were used to determine significant differences. mean ± S.E., \**p* ≤ 0.05, \*\**p* ≤ 0.01, and *n.s*., not significant.

**Supplementary Table 1.**
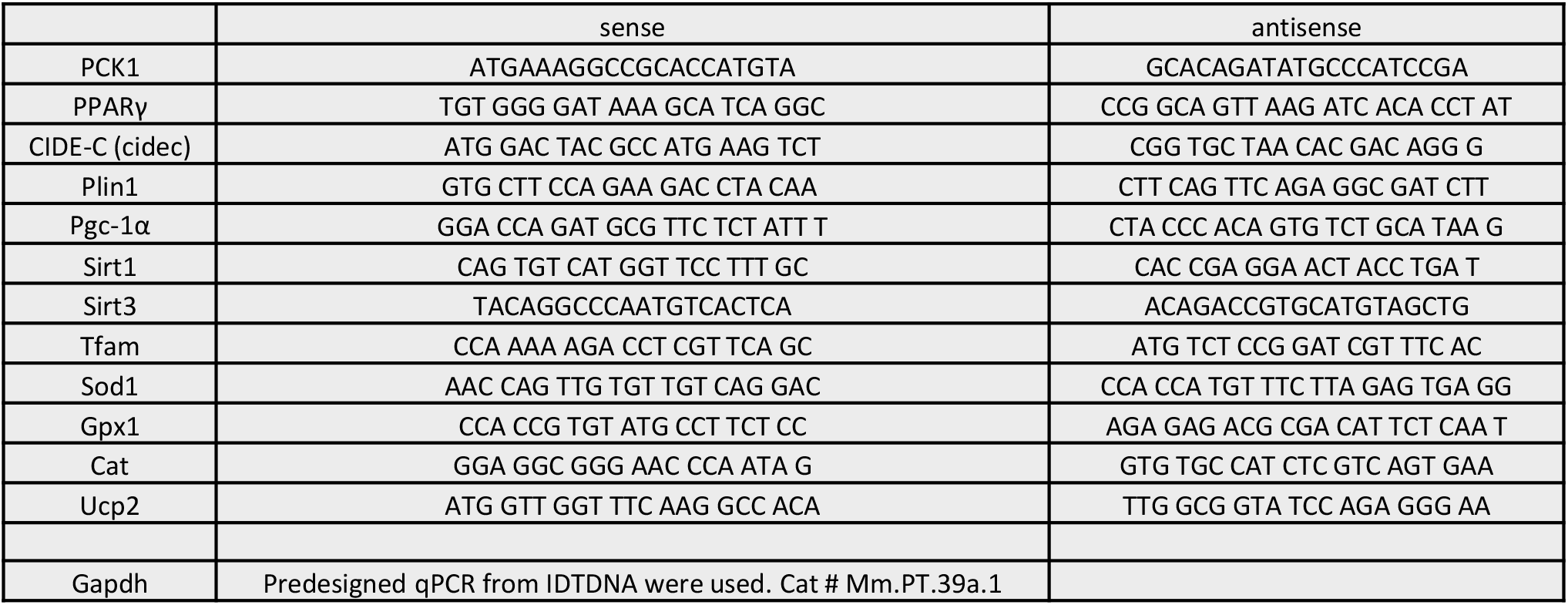
The primers and their sequences that are used for qPCR analysis are listed.

## REFERENCES

1. S. E. Briggs, Special Populations in Implantable Auditory Devices: Geriatric. Otolaryngol. Clin. North Am. 52, 331–339 (2019).

2. J. Wang, J. L. Puel, Presbycusis: An Update on Cochlear Mechanisms and Therapies. J. Clin. Med. 9 (2020), doi:10.3390/JCM9010218.

3. Y. Hou, Y. Wei, S. Lautrup, B. Yang, Y. Wang, S. Cordonnier, M. P. Mattson, D. L. Croteau, V. A. Bohr, NAD+ supplementation reduces neuroinflammation and cell senescence in a transgenic mouse model of Alzheimer’s disease via cGAS-STING. Proc. Natl. Acad. Sci. U. S. A. 118 (2021), doi:10.1073/PNAS.2011226118/SUPPL_FILE/PNAS.2011226118.SAPP.PDF.

4. E. F. Fang, H. Kassahun, D. L. Croteau, M. Scheibye-Knudsen, K. Marosi, H. Lu, R. A. Shamanna, S. Kalyanasundaram, R. C. Bollineni, M. A. Wilson, W. B. Iser, B. N. Wollman, M. Morevati, J. Li, J. S. Kerr, Q. Lu, T. B. Waltz, J. Tian, D. A. Sinclair, M. P. Mattson, H. Nilsen, V. A. Bohr, NAD + Replenishment Improves Lifespan and Healthspan in Ataxia Telangiectasia Models via Mitophagy and DNA Repair. Cell Metab. 24, 566–581 (2016).

5. E. F. Fang, M. Scheibye-Knudsen, L. E. Brace, H. Kassahun, T. Sengupta, H. Nilsen, J. R. Mitchell, D. L. Croteau, V. A. Bohr, Defective mitophagy in XPA via PARP-1 hyperactivation and NAD(+)/SIRT1 reduction. Cell 157, 882–896 (2014).

6. E. Verdin, NAD + in aging, metabolism, and neurodegeneration (; https://www.science.org).

7. K. D. Brown, S. Maqsood, J. Y. Huang, Y. Pan, W. Harkcom, W. Li, A. Sauve, E. Verdin, S. R. Jaffrey, Activation of SIRT3 by the NAD+ precursor nicotinamide riboside protects from noise-induced hearing loss. Cell Metab. 20, 1059–1068 (2014).

8. M. N. Okur, B. Mao, R. Kimura, S. Haraczy, T. Fitzgerald, K. Edwards-Hollingsworth, J. Tian, W. Osmani, D. L. Croteau, M. W. Kelley, V. A. Bohr, Short-term NAD + supplementation prevents hearing loss in mouse models of Cockayne syndrome., doi:10.1038/s41514-019-0040-z.

9. J. Yoshino, J. A. Baur, S. ichiro Imai, NAD+ Intermediates: The Biology and Therapeutic Potential of NMN and NR. Cell Metab. 27, 513–528 (2018).

10. N. Diguet, S. A. J. Trammell, C. Tannous, R. Deloux, J. Piquereau, N. Mougenot, A. Gouge, M. Gressette, B. Manoury, J. Blanc, M. Breton, J. F. Decaux, G. G. Lavery, I. Baczkó, J. Zoll, A. Garnier, Z. Li, C. Brenner, M. Mericskay, Nicotinamide riboside preserves cardiac function in a mouse model of dilated cardiomyopathy. Circulation 137, 2256 (2018).

11. Y. Hou, S. Lautrup, S. Cordonnier, Y. Wang, D. L. Croteau, E. Zavala, Y. Zhang, K. Moritoh, J. F. O’Connell, B. A. Baptiste, T. V. Stevnsner, M. P. Mattson, V. A. Bohr, NAD+ supplementation normalizes key Alzheimer’s features and DNA damage responses in a new AD mouse model with introduced DNA repair deficiency. Proc. Natl. Acad. Sci. U. S. A. 115, E1876–E1885 (2018).

12. C. R. Martens, B. A. Denman, M. R. Mazzo, M. L. Armstrong, N. Reisdorph, M. B. Mcqueen, M. Chonchol, D. R. Seals, Chronic nicotinamide riboside supplementation is well-tolerated and elevates NAD + in healthy middle-aged and older adults., doi:10.1038/s41467-018-03421-7.

13. N. Braidy, G. J. Guillemin, H. Mansour, T. Chan-Ling, A. Poljak, R. Grant, Age related changes in NAD+ metabolism oxidative stress and Sirt1 activity in wistar rats. PLoS One 6 (2011), doi:10.1371/JOURNAL.PONE.0019194.

14. E. F. Fang, Y. Hou, S. Lautrup, M. B. Jensen, B. Yang, T. SenGupta, D. Caponio, R. Khezri, T. G. Demarest, Y. Aman, D. Figueroa, M. Morevati, H. J. Lee, H. Kato, H. Kassahun, J. H. Lee, D. Filippelli, M. N. Okur, A. Mangerich, D. L. Croteau, Y. Maezawa, C. A. Lyssiotis, J. Tao, K. Yokote, T. E. Rusten, M. P. Mattson, H. Jasper, H. Nilsen, V. A. Bohr, NAD+ augmentation restores mitophagy and limits accelerated aging in Werner syndrome. Nat. Commun. 2019 101 10, 1–18 (2019).

15. M. N. Okur, E. F. Fang, E. M. Fivenson, V. Tiwari, D. L. Croteau, V. A. Bohr, Cockayne syndrome proteins CSA and CSB maintain mitochondrial homeostasis through NAD+ signaling. Aging Cell 19 (2020), doi:10.1111/acel.13268.

16. D. Malide, J. Liu, I. I. Rovira, C. A. Combs, N. Sun, T. Finkel, A fluorescence-based imaging method to measure in vitro and in vivo mitophagy using mt-Keima. Nat. Protoc. 12 (2016), doi:10.1038/nprot.2017.060.

17. Y. Sergeyenko, K. Lall, M. Charles Liberman, S. G. Kujawa, Age-related cochlear synaptopathy: an early-onset contributor to auditory functional decline. J. Neurosci. 33, 13686–13694 (2013).

18. B. Grygiel-Górniak, Peroxisome proliferator-activated receptors and their ligands: Nutritional and clinical implications - A review. Nutr. J. 13, 1–10 (2014).

19. M. Sekulic-Jablanovic, V. Petkovic, M. B. Wright, K. Kucharava, N. Huerzeler, S. Levano, Y. Brand, K. Leitmeyer, A. Glutz, A. Bausch, D. Bodmer, Effects of peroxisome proliferator activated receptors (PPAR)-γ and -α agonists on cochlear protection from oxidative stress. 12 (2017), doi:10.1371/JOURNAL.PONE.0188596.

20. A. Gorga, G. M. Rindone, & M. Regueira, E. H. Pellizzari, M. C. Camberos, S. B. Cigorraga, M. F. Riera, & M. N. Galardo, S. B. Meroni, PPARγ activation regulates lipid droplet formation and lactate production in rat Sertoli cells., doi:10.1007/s00441-017-2615-y.

21. Z. Sun, J. Gong, H. Wu, W. Xu, L. Wu, D. Xu, J. Gao, J. W. Wu, H. Yang, M. Yang, P. Li, Perilipin1 promotes unilocular lipid droplet formation through the activation of Fsp27 in adipocytes. Nat. Commun. 4 (2013), doi:10.1038/NCOMMS2581.

22. T. H. M. Grahn, Y. Zhang, M. J. Lee, A. G. Sommer, G. Mostoslavsky, S. K. Fried, A. S. Greenberg, V. Puri, FSP27 and PLIN1 interaction promotes the formation of large lipid droplets in human adipocytes. Biochem. Biophys. Res. Commun. 432, 296–301 (2013).

23. D. Xu, Z. Wang, Y. Xia, F. Shao, W. Xia, Y. Wei, X. Li, X. Qian, J. H. Lee, L. Du, Y. Zheng, G. Lv, J. shiun Leu, H. Wang, D. Xing, T. Liang, M. C. Hung, Z. Lu, The gluconeogenic enzyme PCK1 phosphorylates INSIG1/2 for lipogenesis. Nat. 2020 5807804 580, 530–535 (2020).

24. R. A. Urrutia, F. Kalinec, Biology and Pathobiology of Lipid Droplets and their Potential Role in the Protection of the Organ of Corti. Hear. Res. 330, 26 (2015).

25. K. Kuramoto, T. Okamura, T. Yamaguchi, T. Y. Nakamura, S. Wakabayashi, H. Morinaga, M. Nomura, T. Yanase, K. Otsu, N. Usuda, S. Matsumura, K. Inoue, T. Fushiki, Y. Kojima, T. Hashimoto, F. Sakai, F. Hirose, T. Osumi, Perilipin 5, a lipid droplet-binding protein, protects heart from oxidative burden by sequestering fatty acid from excessive oxidation. J. Biol. Chem. 287, 23852–23863 (2012).

26. M. A. Aon, N. Bhatt, S. Cortassa, Mitochondrial and cellular mechanisms for managing lipid excess. Front. Physiol. 5 (2014), doi:10.3389/FPHYS.2014.00282.

27. A. P. Bailey, G. Koster, C. Guillermier, E. M. A. Hirst, J. I. MacRae, C. P. Lechene, A. D. Postle, A. P. Gould, Antioxidant Role for Lipid Droplets in a Stem Cell Niche of Drosophila. Cell 163, 340–353 (2015).

28. S. H. Sha, A. Kanicki, G. Dootz, A. E. Talaska, K. Halsey, D. Dolan, R. Altschuler, J. Schacht, Age-related auditory pathology in the CBA/J mouse. Hear. Res. 243, 87 (2008).

29. S. G. Kujawa, M. C. Liberman, Synaptopathy in the noise-exposed and aging cochlea: Primary neural degeneration in acquired sensorineural hearing loss. Hear. Res. 330, 191–199 (2015).

30. S. Safieddine, A. El-Amraoui, C. Petit, The Auditory Hair Cell Ribbon Synapse: From Assembly to Function. (2012), doi:10.1146/annurev-neuro-061010-113705.

31. V. G. Magupalli, K. Schwarz, K. Alpadi, S. Natarajan, G. M. Seigel, F. Schmitz, Multiple RIBEYE-RIBEYE interactions create a dynamic scaffold for the formation of synaptic ribbons. J. Neurosci. 28, 7954–7967 (2008).

32. J. A. Olzmann, P. Carvalho, Dynamics and functions of lipid droplets. Nat. Rev. Mol. Cell Biol., doi:10.1038/s41580-018-0085-z.

33. J. S. Hansen, S. De Maré, H. A. Jones, O. Göransson, K. Lindkvist-Petersson, Visualization of lipid directed dynamics of perilipin 1 in human primary adipocytes. Sci. Reports 2017 71 7, 1–14 (2017).

34. M. Boutant, S. S. Kulkarni, M. Joffraud, J. Ratajczak, M. Valera-Alberni, R. Combe, A. Zorzano, C. Cantó, M. Valera-Alberni, R. Combe, A. Zorzano, C. Cantó, Mfn2 is critical for brown adipose tissue thermogenic function. EMBO J. 36 (2017), doi:10.15252/EMBJ.201694914.

35. G. Sharma, R. Saubouny, M. M. Joel, K. Martens, D. Martino, A. P. J. de Koning, G. Pfeffer, T. E. Shutt, Characterization of a novel variant in the HR1 domain of MFN2 in a patient with ataxia, optic atrophy and sensorineural hearing loss. bioRxiv, 2021.01.11.426268 (2021).

36. M. Shimizu, A. Takeshita, T. Tsukamoto, F. J. Gonzalez, T. Osumi, Tissue-selective, bidirectional regulation of PEX11 alpha and perilipin genes through a common peroxisome proliferator response element. Mol. Cell. Biol. 24, 1313–1323 (2004).

37. K. Okabe, A. Nawaz, Y. Nishida, K. Yaku, I. Usui, K. Tobe, T. Nakagawa, NAD+ Metabolism Regulates Preadipocyte Differentiation by Enhancing α-Ketoglutarate-Mediated Histone H3K9 Demethylation at the PPARγ Promoter. Front. Cell Dev. Biol. 8, 1409 (2020).

38. S. L. Johnson, A. Forge, M. Knipper, S. Münkner, W. Marcotti, Tonotopic Variation in the Calcium Dependence of Neurotransmitter Release and Vesicle Pool Replenishment at Mammalian Auditory Ribbon Synapses. J. Neurosci. 28, 7670–7678 (2008).

39. S. Han, Z. Du, K. Liu, S. Gong, Nicotinamide riboside protects noise-induced hearing loss by recovering the hair cell ribbon synapses. Neurosci. Lett. 725, 134910 (2020).

40. Y. L. M. Ying, C. D. Balaban, Regional distribution of manganese superoxide dismutase 2 (Mn SOD2) expression in rodent and primate spiral ganglion cells. Hear. Res. 253, 116–124 (2009).

41. C. Cantó, R. H. Houtkooper, E. Pirinen, D. Y. Youn, M. H. Oosterveer, Y. Cen, P. J. Fernandez-Marcos, H. Yamamoto, P. A. Andreux, P. Cettour-Rose, K. Gademann, C. Rinsch, K. Schoonjans, A. A. Sauve, J. Auwerx, The NAD+ precursor nicotinamide riboside enhances oxidative metabolism and protects against high-fat diet induced obesity. Cell Metab. 15, 838 (2012).

42. M. Shiozaki, N. Hayakawa, M. Shibata, M. Koike, Y. Uchiyama, T. Gotow, Closer association of mitochondria with lipid droplets in hepatocytes and activation of KupVer cells in resveratrol-treated senescence-accelerated mice. Histochem. Cell Biol. 136, 475–489 (2011).

43. G. M. Kalinec, C. Park, P. Thein, F. M. Kalinec, Working with Auditory HEI-OC1 Cells. J. Vis. Exp, 54425 (2016).

44. K. Vikhe Patil, B. Canlon, C. R. Cederroth, High quality RNA extraction of the mammalian cochlea for qRT-PCR and transcriptome analyses. Hear. Res. 325, 42–48 (2015).

45. S. Anders, W. Huber, Differential expression analysis for sequence count data. Genome Biol. 11, 1–12 (2010).

46. M. Müller, K. Von Hünerbein, S. Hoidis, J. W. T. Smolders, A physiological place-frequency map of the cochlea in the CBA/J mouse. Hear. Res. 202, 63–73 (2005).

47. E. F. Fang, K. Palikaras, N. Sun, E. M. Fivenson, R. D. Spangler, J. S. Kerr, S. A. Cordonnier, Y. Hou, E. Dombi, H. Kassahun, N. Tavernarakis, J. Poulton, H. Nilsen, V. A. Bohr, In Vitro and In Vivo Detection of Mitophagy in Human Cells, C. Elegans, and Mice. J. Vis. Exp. 2017, 56301 (2017).

